# A unifying framework disentangles genetic, epigenetic, and stochastic sources of drug-response variability in an *in vitro* model of tumor heterogeneity

**DOI:** 10.1101/2020.06.05.136119

**Authors:** Corey E. Hayford, Darren R. Tyson, C. Jack Robbins, Peter L. Frick, Vito Quaranta, Leonard A. Harris

**Affiliations:** Chemical and Physical Biology Graduate Program, Vanderbilt University School of Medicine, Nashville, TN USA; Department of Biochemistry, Vanderbilt University School of Medicine, Nashville, TN USA; Department of Pharmacology, Vanderbilt University School of Medicine, Nashville, TN USA

**Author notes:** MD-PhD Graduate Program, Yale University School of Medicine, New Haven, CT USA.

## Abstract

Tumor heterogeneity is a primary cause of treatment failure and acquired resistance in cancer patients. Even in cancers driven by a single mutated oncogene, variability of targeted therapy response is observed. Additional genetic mutations can only partially explain this variability, leading to consideration of non-genetic factors, such as “stem-like” and “mesenchymal” phenotypic states, as critical contributors to tumor relapse and resistance. Here, we show that both genetic and non-genetic factors contribute to targeted drug-response variability in an experimental tumor heterogeneity model based on multiple versions and clonal sublines of PC9, the archetypal EGFR-mutant non-small cell lung cancer cell line. We observe significant drug-response variability across PC9 cell line versions, among sublines, and within sublines. To disentangle genetic, epigenetic, and stochastic components underlying this variability, we adopt a theoretical framework whereby distinct genetic states give rise to multiple epigenetic “basins of attraction”, across which cells can transition driven by stochastic factors such as gene expression noise and asymmetric cell division. Using mutational impact analysis, single-cell differential gene expression, and semantic similarity of gene ontology terms to connect genomics and transcriptomics, we establish a baseline of genetic differences explaining drug-response variability across PC9 cell line versions. In contrast, with the same approach, we conclude that in all but one of the clonal sublines, drug-response variability is due to epigenetic rather than genetic differences. Finally, using a clonal drug-response assay and stochastic simulations, we attribute drug-response variability within sublines to intracellular stochastic fluctuations and confirm that one subline likely contains a genetic resistance mutation that emerged in the absence of selective pressures. We propose that a theoretical framework deconvolving the complex interplay among genetic, epigenetic, and stochastic sources of intratumoral heterogeneity will lead to novel therapeutic strategies to combat tumor relapse and resistance.

## INTRODUCTION

Cancer is a complex and dynamic disease characterized by intertumoral and intratumoral heterogeneities that have been implicated in treatment avoidance and acquired resistance to therapy^1, 2^. Genetic differences among cancer cells within and across tumors have long been appreciated^3–8^. Indeed, genomic instability is a hallmark of cancer^9, 10^ and is considered to be the primary source of this genetic heterogeneity^11, 12^. However, it is becoming increasingly apparent that genetics alone cannot fully explain the wide ranges of responses observed in patient populations to anticancer therapies^13, 14^. Researchers are, therefore, increasingly looking to *non-genetic* sources of tumor heterogeneity for explanations. Broadly speaking, non-genetic heterogeneity comes in two forms:^13, 15–19^ *epigenetic*, which is heritable^15, 17^ (at least for a few generations), and *stochastic*, which is not heritable and is due to intrinsic factors such as noise in gene expression^20–24^ and asymmetric cell division^25, 26^. Non-genetic heterogeneity has been linked to drug tolerance and decreases in drug sensitivity *in vitro*^2, 27–30^, *in vivo*^27, 28, 31^, and clinically^32, 33^.

A theoretical framework for understanding the connections between genetic and non-genetic sources of tumor heterogeneity is the “epigenetic landscape,” which was proposed by Waddington over 50 years ago^34^ but has received renewed attention recently^16, 17, 35, 36^. In analogy to the potential energy landscape of physical chemistry, Waddington posited that a cellular state can be assigned a “quasi-potential” energy and placed within a landscape where local minima correspond to cellular phenotypes. Phenotypic state transitions occur when cells traverse the barriers separating adjacent basins, driven by intrinsic or extrinsic sources of noise^37, 38^. Importantly, the topography of the epigenetic landscape depends on a complex set of biochemical interactions within (and possibly spanning^39^) cells and the values of associated rate parameters^40, 41^. Many of these parameters depend strongly on protein structure (e.g., the accessibility of a binding domain), which is encoded within the DNA. Thus, an epigenetic landscape can be thought of as deriving from a particular “genetic state”^16, 36^ (see below). Typical tumors comprise numerous genetic states^6, 42^ and, thus, in this framework have numerous overlapping epigenetic landscapes, each of which is subject to noise-induced phenotypic transitions (FIG. 1A). Over time, the composition of a tumor is expected to evolve through a series of genetic mutations and expansions, as in the standard picture of clonal evolution^8, 17^, but with the important distinction that each genetic clone comprises numerous epigenetic states (FIG. 1B), among which cells can interconvert and some of which may be well suited to survive future stressors, such as drug treatments^30, 43^.

**Figure 1.**
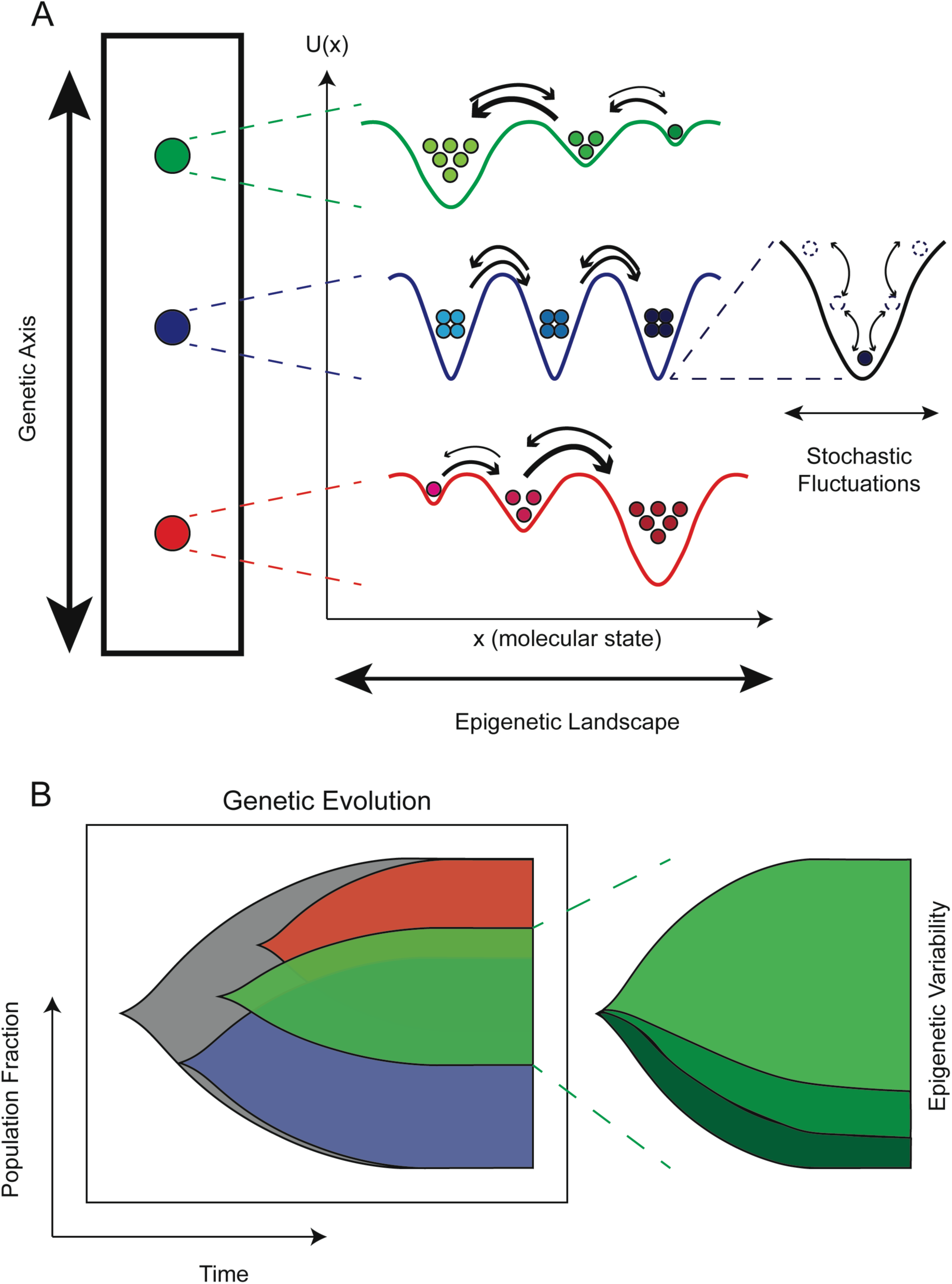
Conceptual G/E/S framework of tumor heterogeneity. (A) Multiple levels of heterogeneity operate within tumors. (*left*) The genetic (G) “axis” defines mutational differences that have an effect on phenotype (e.g., drug sensitivity). (*middle*) Each genetic clone has an associated epigenetic (E) landscape, where cells are distributed amongst basins, known as attractors. The topography of the epigenetic landscape is defined by the dynamical biochemical network that controls cell fate and function. Molecular state (defined as *x*) is the x-axis and quasi-potential energy (defined as *U(x)*) is the y-axis. (*right*) Cell states experience stochastic (S) fluctuations within epigenetic basins. Most fluctuations are minor and do not significantly change the cell state but occasionally a large fluctuation results in a barrier crossing, leading to a phenotypic state change. (B) Schematic view of genetic clonal evolution, where clones initiate and expand from a recent common ancestor (grey) at different points in time during progression. The evolution of a single genetic clone (*right*) is highlighted to illustrate that cells populate different epigenetic states during expansion, some of which may be drug tolerant.

This “systems” view of tumor heterogeneity, as a three-tiered amalgamation of genetic, epigenetic, and stochastic sources of variability, has been proposed previously^16, 17, 35, 36^ to explain the wide ranges of responses to, and failures of, anticancer therapies within genomically stratified patient populations^44, 45^. For instance, epidermal growth factor receptor (EGFR) inhibitors are not equally effective across patients with EGFR-mutant lung cancer and in almost all cases tumors eventually acquire resistance^46, 47^. However, without strong experimental evidence to support it, this framework has yet to achieve broad acceptance within the cancer research community^36^. Here, we tackle this issue by performing extensive drug-response profiling, followed by genomic and transcriptomic characterization and mathematical population dynamics modeling, of multiple “versions” and single cell-derived sublines of an archetypal non-small cell lung cancer (NSCLC) cell line. We chose this family of cell line versions and sublines because we suspected, based on the individual lineages and the conditions under which each member was derived, that it would effectively mimic the constitution of a genetically and epigenetically heterogeneous tumor (see Supplementary Note for further discussion). The cell line versions include two variants that have been maintained independently for many years at separate institutions and one (derived from one of these variants) with a known genetic resistance mutation acquired through systematic dose escalation. Single cell-derived sublines, isolated from one of these versions and passaged numerous times prior to experimental interrogation, model epigenetic variation within an individual genetic state. We use bulk genomic and single-cell RNA sequencing of the cell line versions to provide a benchmark for genetic variability (differences across populations; see Supplementary Table S1) against which we compare genomic and transcriptomic differences seen among the sublines. Our analysis suggests that all but one of the sublines are genetically identical but differ epigenetically, i.e., they constitute distinct basins within a common epigenetic landscape. The monoclonality of these sublines is further supported by stochastic simulations that indicate drug response variability seen across isolated colonies can be explained by intrinsic randomness in cell division and death. We also detail a case where one subline appears to contain at least two distinct genetic states, the second presumably acquired after establishment of the subline. Below, we provide a detailed description of the genetic-epigenetic-stochastic (G/E/S) framework on which this work is based, with precise definitions of relevant terms and concepts, followed by experimental and theoretical analyses at the genomic, transcriptomic, and cell population levels. We conclude with a discussion of the consequences of this view of tumor heterogeneity for how we think about cancer and for devising effective anticancer strategies in the future.

## RESULTS

### A theoretical framework for deconvolving intratumoral heterogeneity into genetic, epigenetic, and stochastic components

Based on prior work in this area^8, 16, 17, 19, 36, 41, 48^, we view intratumoral heterogeneity as comprised of genetic, epigenetic, and stochastic components (FIG. 1A) that are broadly distinguished based on characteristic timescale of change: functional genetic mutations are acquired on the order of weeks to months^49, 50^, transitions between epigenetic states occur on the order of hours to weeks^51^, and stochastic fluctuations in protein concentrations, and other sources of intracellular noise, operate on the order of seconds to minutes^52^. Separating processes based on characteristic timescale is a common approach in physics^53^ and has a long history in biology^54^. Note that we distinguish between “heterogeneity”, defined as differences (genetic, epigenetic, or stochastic) that exist *within* a cell population, and “variability”, which are differences seen *across* populations. A tumor is heterogeneous if it is composed of subpopulations that display variabilities amongst each other. Below, we describe in detail each of these levels of heterogeneity, with precise definitions of terms that will be important for later analyses (see also Supplementary Table S1).

#### Genetic level (FIG. 1A, left)

Genomic instability is a hallmark of cancer^9, 10^. For cancer cells in culture, it has been estimated that approximately five million cell divisions are necessary to acquire at least one mutation per gene^49^. With more than 20,000 genes in the genome, this amounts to one mutation every two weeks on average. These mutations can be single-nucleotide polymorphisms (SNPs), insertions/deletions (InDels), and larger structural variations (see Supplementary Note and Supplementary FIG. S1). While mutations in the genome are relatively frequent, many do not have functional consequence, so-called “passenger” mutations^16^. Therefore, we make a distinction between the *genomic state* of a cell and the *genetic state*: the genomic state is the full sequence of nucleotides that make up the DNA whereas the genetic state is the subset of this sequence that contributes to a specific cellular phenotype^55, 56^. In other words, *two cells may differ genomically but be genetically identical if the mutations that differentiate them occur in portions of the genome that have no functional consequence on the phenotype of interest* (see Supplementary FIG. S1A). As such, the timescale for generating genetic heterogeneity within a population of cancer cells can be quite slow (much slower than the acquisition of genomic heterogeneity), on the order of weeks to months^49, 50^. Note that enumerating the subset of the genome that defines the genetic state can be difficult (if not impossible) in practice and depends on the phenotypic context^57^. Here, we use drug response as the phenotype of interest. Thus, the genetic state is, in principle, defined over all genes associated with drug response, which includes those involved in stress response, metabolism, cell cycle regulation, and many others.

#### Epigenetic level (FIG. 1A, middle)

Conceptualized as a quasi-potential energy surface where local minima, or “basins of attraction,” correspond to cellular phenotypes, epigenetic landscapes were first proposed by Waddington as an abstract tool for understanding phenotypic plasticity and cellular differentiation during development^34^. The genetic state of a cell sets the topography of the landscape^8, 36^ and intrinsic (e.g., gene expression^20^) or extrinsic^14, 58^ sources of noise drive transitions between phenotypes. In cancer, rather than a well-defined hierarchy, multiple epigenetic states of comparable stability can coexist^59, 60^, often with differential drug sensitivities^27, 61, 62^. A population of cancer cells would thus be expected to spread out across these phenotypes, producing a highly heterogeneous mix, even among isogenic cells^59^. Phenotypic state transitions have been observed to occur on the order of hours to weeks^30, 62^, meaning epigenetic diversification occurs on a much faster timescale than genetic diversification. From a molecular perspective, the epigenetic landscape is the consequence of the complex biochemical interaction networks underlying cellular function^63, 64^. Complex dynamical networks can harbor multiple stable states, termed “attractors”, towards which a system will tend to return (relax back) in response to small perturbations^65, 66^. This property underlies epigenetic heritability: daughter cells inherit similar molecular contents as their parent; hence, they tend to remain within the region of influence of the parental attractor. Here, we use the transcriptional state of a cell, revealed through single-cell transcriptomics, as a proxy for the epigenetic state, with the understanding that mRNA is only one lens through which to view epigenetics. Indeed, this definition of an epigenetic state, i.e., as a stable state of a complex biochemical network, differs from the traditional molecular biology definition in terms of “epigenetic marks”^67^ (e.g., DNA methylation, regions of open chromatin). The two are related, however, in the sense that the same biochemical network that defines the epigenetic landscape also sets the epigenetic marks on the DNA^19^ (see Supplementary Note for an in depth discussion).

#### Stochastic level (FIG. 1A, right)

Fluctuations, or noise, in intracellular species concentrations have long been recognized as a source of non-genetic heterogeneity in isogenic cell populations, first in bacteria^52, 68^ and then in yeast^69, 70^ and mammalian cells^59, 62, 71^. Intracellular noise can be “intrinsic”, i.e., due to the probabilistic nature of biochemical interactions^72, 73^, or “extrinsic”^38^, affecting the *rates* of interactions, synthesis, and degradation within a biochemical network. Intrinsic sources of noise include transcriptional bursting^74, 75^, translational bursting^76^, and randomness in mRNA/protein degradation^77^, oligomerization^78^, and post-translational modification^79^. Extrinsic noise includes randomness in the distribution of molecular contents upon cell division^26, 80^; environmental factors such as inhomogeneities in cell culture media^81^, fluctuations in temperature^82^ and pH^83^, and spatial variations in the microenvironment^84^. Importantly, fluctuations at the molecular level can drive probabilistic cell fate decisions^20, 21, 23, 85–87^, including division and death^88, 89^, at the single-cell level and phenotypic diversification at the epigenetic (population) level^59, 90^. Experimentally, at the intracellular level, intrinsic and extrinsic noise are difficult to distinguish, generally requiring multiple fluorescent reporters and the ability to fine-tune the external environment^38^. Theoretically, the chemical Master Equation^91, 92^ (CME) is the construct upon which all stochastic dynamical analyses are based. Since the CME is difficult to solve in general, a number of stochastic algorithms have been developed for simulating fluctuations at both the single cell and cell population levels^73, 93^.

### Cell line versions and single cell-derived sublines exhibit drug-response variability at the cell population level

We chose canonical NSCLC cell line PC9^94^ as the model system to substantiate the G/E/S heterogeneity framework described above. The PC9 cell line is characterized by an EGFR-ex19del mutation (Supplementary FIG. S2A), making it sensitive to inhibition of the mutant EGFR protein. We utilized three versions of the cell line: PC9-VU, originating from Vanderbilt University^95^; PC9-MGH, maintained at Massachusetts General Hospital^62, 96^; and PC9-BR1, derived from PC9-VU and containing a known secondary resistance mutation (EGFR-T790M) obtained through dose escalation therapy in the EGFR inhibitor (EGFRi) afatinib^95^. Although it is unclear when the PC9-VU and PC9-MGH versions, which originate from a common founder cell population, were independently established (see Supplementary FIG. S2B), both maintain the oncogenic mutation in the EGFR gene and show sensitivity to EGFR inhibition^62, 89^. In the absence of drug, PC9-VU and PC9-BR1 have essentially identical proliferation rates, while PC9-MGH grows at a slightly lower rate (Supplementary FIG. S3A). However, in response to the EGFRi erlotinib (see Methods), the three cell line versions display drastically different drug sensitivities (FIG. 2A): PC9-MGH exhibits substantial cell death after an initial “drug stabilization” phase (∼72h), PC9-VU settles into a near-zero rate of growth, and PC9-BR1 displays insensitivity to EGFRi, as expected. These results are consistent with the high sensitivity of PC9-MGH to erlotinib reported in Sharma et al.^62^ and the lower sensitivity of PC9-VU observed in Harris et al.^88, 89^

**Figure 2.**
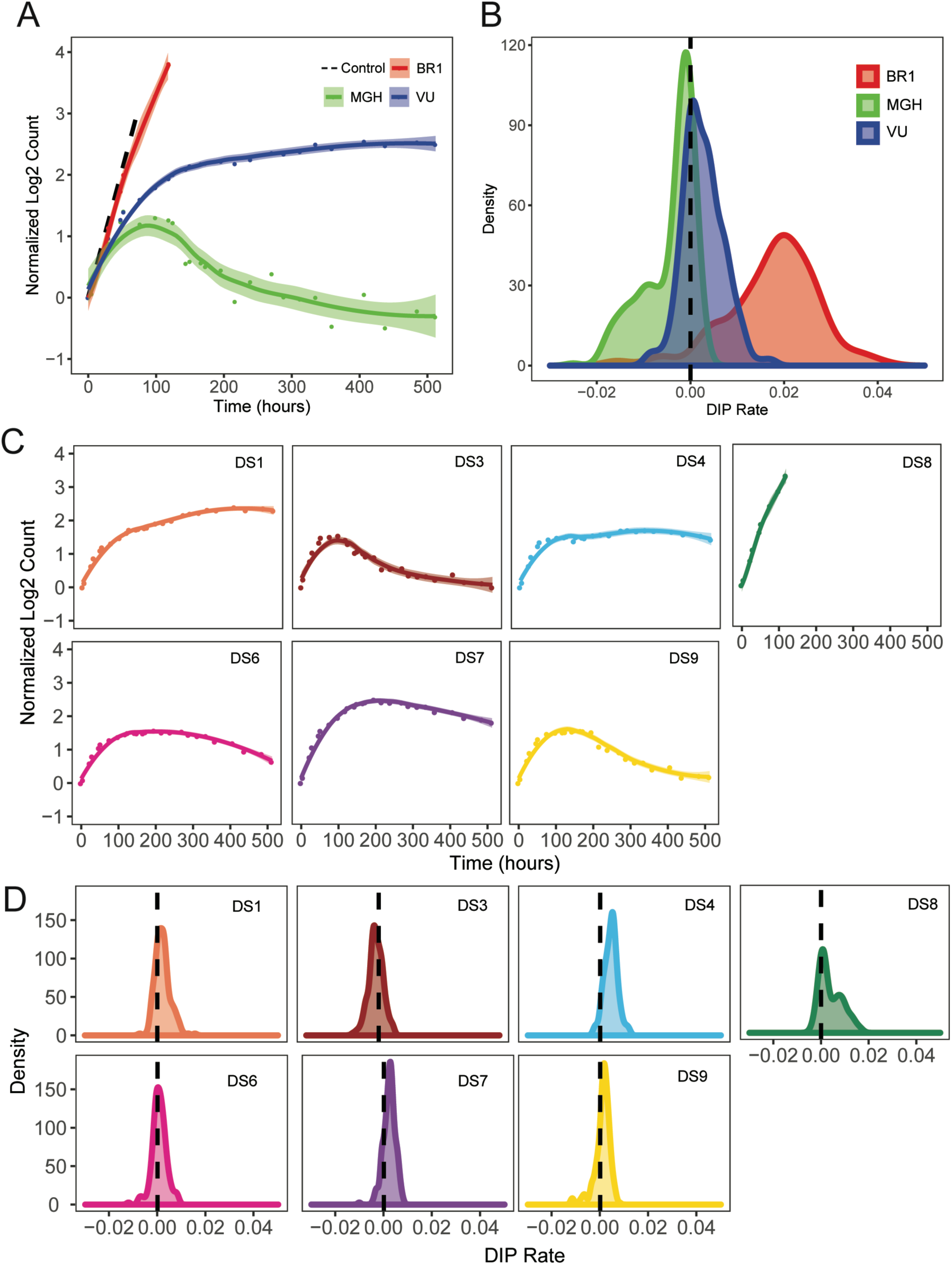
Phenotypic differences among PC9 cell line versions and discrete sublines quantified in terms of drug response. (A) Population growth curves for three cell line versions treated with 3μM erlotinib for approximately three weeks, plus vehicle control. (B) Drug-induced proliferation (DIP) rate distributions compiled from single-colony growth trajectories under erlotinib treatment (3uM) in a cFP assay. DIP rates are calculated from the growth curves 48h post-drug addition to the end of the experiment. (C) Seven discrete sublines (DS), derived from PC9-VU, were treated with 3μM erlotinib for three weeks. (D) DIP rate distributions from a cFP assay of the sublines in 3μM erlotinib. In *A* and *C*, dots are the means of six experimental replicates at each time point; solid lines are best fits to the drug response trajectories with point-wise 95% confidence intervals. In *B* and *D*, DIP rate distributions are plotted as kernel density estimates; dashed black lines signify zero DIP rate, for visual orientation.

We also quantified clonal drug response variability using clonal Fractional Proliferation^97^ (cFP), an assay that tracks the growth of many single cell-derived colonies over time and quantifies drug sensitivity for each colony in terms of the drug-induced proliferation (DIP) rate^88, 89^, defined as the stable rate of proliferation achieved after extended drug exposure (see Methods and Supplementary Note). We performed cFP on each cell line version treated with erlotinib and observed wide ranges of drug responses across colonies (Supplementary FIG. S3B) and substantial differences in the aggregate responses across versions (FIG. 2B). The distribution of DIP rates for PC9-BR1 lies almost entirely in the positive DIP rate range and is clearly distinct from the others. The PC9-VU and PC9-MGH distributions have significant overlap but the PC9-MGH distribution has a marked shoulder in the negative DIP rate region while the PC9-VU distribution extends further into the positive range. These distributions are consistent with and explain the differential drug responses observed between the cell line versions (FIG. 2A): PC9-BR1 is resistant to EGFRi because its DIP rate distribution is entirely in the positive range, PC9-VU goes into a near-zero (slightly positive) growth phase because its DIP rate distribution is centered near zero, and the large shoulder in the PC9-MGH distribution explains why it exhibits significant cell death in the period immediately following drug treatment.

In addition, several single cell-derived discrete sublines (DS1, DS3, DS4, DS6, DS7, DS8, DS9) were isolated from PC9-VU and subjected to the same analyses as above. In the absence of drug, all sublines grow at almost equal rates in culture (Supplementary FIG. S3C). However, in the presence of EGFRi, the sublines exhibit a wide range of responses, from positive growth to negative (FIG. 2C). When overlaid with the cFP results for PC9-VU, we see that the subline responses broadly recapitulate the observed variability seen in the parental line (Supplementary FIG. S4A). A notable exception is DS8, which is essentially resistant to EGFRi, having only a slightly lower proliferation rate than the fully resistant PC9-BR1 (Supplementary FIG. S4B). We also performed cFP assays on the sublines under erlotinib treatment (FIG. 2D). Interestingly, similar to the cell line versions, we found that the sublines also exhibit distributions of DIP rates, albeit generally narrower than those for the versions. The subline distributions have a large degree of overlap with one another but the medians of the distributions are statistically distinct (Supplementary FIG. S4C). DS8 is again an exception, exhibiting a bimodal DIP rate distribution with a major mode centered close to zero and a large shoulder in the positive DIP rate region.

### Cell line versions differ significantly at the genetic and transcriptomic levels

Given how long the PC9-VU and PC9-MGH cell line versions have been maintained separately, it is virtually certain that they differ genetically^42^. We also know that PC9-BR1 contains a known genetic resistance mutation and likely numerous additional mutations acquired during dose escalation. Thus, we perform bulk whole exome sequencing (WES) and single-cell RNA sequencing (sc-RNA-seq) on the cell line versions in order to establish a benchmark for genetic variation against which we can compare the sublines.

From WES, we identify mutations (SNPs and InDels) in each cell line version that pass a specified threshold for variant detection (see Methods and Supplementary FIG. S5A,B) and calculate the number of mutations per chromosome (FIG. 3A). We see a large amount of variability in the number of called variants between cell line versions (average coefficient of variation (CV) per chromosome = 12.84). We also identify mutations unique to each cell line version (Supplementary FIG. S5C) and calculate the proportionality of unique mutations compared to the total number of mutations (FIG. 3B). Although a majority of the mutations are shared between versions, a significant number are unique: PC9-BR1 has the largest proportional representation of unique mutations, followed by PC9-MGH and then PC9-VU. Furthermore, we annotate unique mutations within each cell line version with an IMPACT score, calculated using the Variant Effect Predictor^98^ (VEP), which classifies severity of variants based on genetic variant annotation and predicted effects (e.g., amino acid change, protein structure modification; see Methods). The IMPACT score differentiates mutations based on a variety of factors to predict whether a mutation is likely to have a phenotypic effect. Categorizing mutations into *low*, *moderate*, and *high* IMPACT score shows that PC9-BR1 has many more impactful mutations than PC9-MGH and PC9-VU, which have similar numbers to each other (FIG. 3C). However, as a *percentage*, only 1% of PC9-MGH unique mutations are predicted to be impactful, compared to 11% in PC9-VU (FIG. 3C), suggesting that PC9-MGH harbors a large number of passenger mutations.

**Figure 3.**
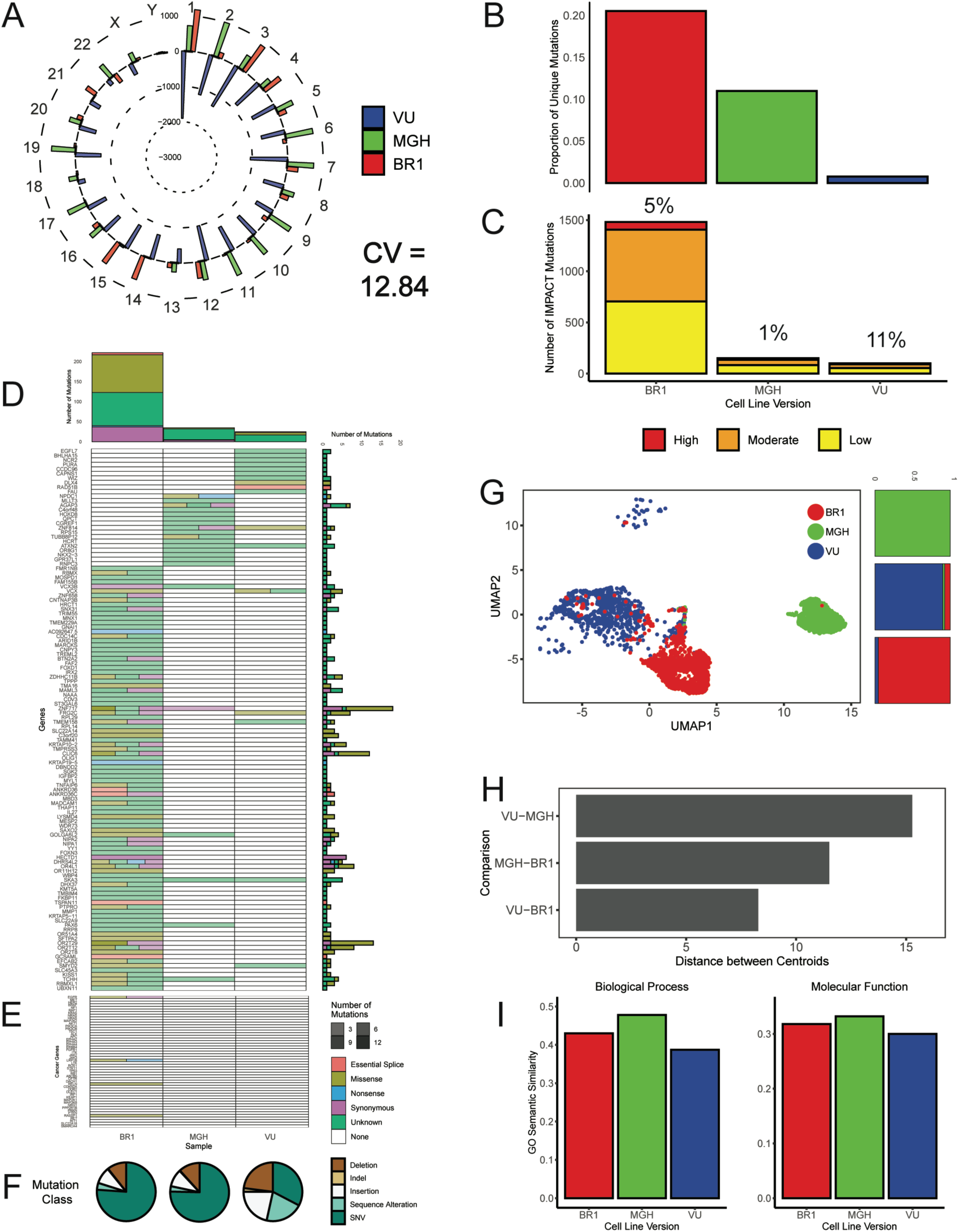
Genomic and transcriptomic characterizations of PC9 cell line versions. (A) Mean-centered mutation count by chromosome for all cell line versions. For each chromosome, versions with fewer mutations than the mean have a bar pointing inwards, while those with more mutations point outwards. Chromosome numbers are noted on the outside edge of the circle. Average coefficient of variation (COV) across all chromosomes is noted. (B) Proportions of unique mutations for all cell line versions. (C) Numbers of IMPACT mutations unique to each cell line version, stratified by IMPACT classification (*low*, *moderate*, *high*). Percentage of unique IMPACT mutations relative to the total number of unique mutations for each cell line version is denoted above each bar. (D) Quantification of mutational differences between cell line versions based on a signature of genes with a high nonsynonymous mutational load. Heatmap elements are colored based on type of mutation: *Essential Splice Site*, *Missense*, *Nonsense*, *Synonymous*, and *Unknown*. Transparency of heatmap elements corresponds to the number of mutations in the gene-mutation type pair (more mutations, less transparent). Total numbers of mutations (stratified by mutation type) across genes and cell line versions are shown as bar plots to the right and above of the heatmap, respectively. (E) Same as *D*, but for a literature-curated set of cancer-associated genes implicated in lung cancer. (F) Mutation class pie charts. All unique mutations in each cell line version were classified as single nucleotide polymorphisms (SNPs), insertions, deletions, indels (combinations of insertions and deletions), or sequence alterations. (G) Uniform Manifold Approximation and Projection (UMAP) visualization of single-cell transcriptomes for cell line versions. For comparison purposes, the UMAP space is defined over all eight PC9 samples (including cell line versions and PC9-VU sublines; see Methods). Proportions of cells in defined clusters (k=3) are shown as bar plots to the right, colored by cell line version (*cluster 1*: 0.1% BR1, 99.9% MGH, 0% VU; *cluster 2*: 7.4% BR1, 2.5% MGH, 90.1% VU; *cluster 3*: 95.5% BR1, 0.1% MGH, 4.4% VU). (H) Pairwise (Euclidian) distances between the centroids of the transcriptional features in *G* for all cell line versions. (I) Gene ontology (GO) semantic similarity analysis of differentially expressed genes (DEGs) and unique IMPACT mutations in each cell line version for “Biological Process” (*left*) and “Molecular Function” (*right*) ontology types.

We also perform a mutational significance analysis on the unique mutations. Based on nonsynonymous mutational load, genes are selected to create a mutational signature of genetic differences within each cell line version and displayed as a heatmap (see Methods). We see that multiple mutations in the signature distinguish the cell line versions (FIG. 3D). Additionally, we generate a literature-curated, cancer-associated gene signature that includes mutations predicted to be implicated in cancer^2, 99^ (see Methods). Only PC9-BR1, which has a known resistance mutation (EGFR-T790M; noted as a missense mutation in the heatmap), harbors any of the mutations in the signature (FIG. 3E). Breakdowns of mutations by type provide additional evidence that PC9-VU mutation representation is distinct and proportionally impactful (FIG. 3F). Together, these genomic data (FIG. 3A–F) starkly illustrate the genetic differences among the cell line versions.

At the transcriptomic level, we use sc-RNA-seq to identify gene expression differences between the cell line versions (see Methods and Supplementary FIG. S6A). After feature selection (Supplementary FIG. S6B), we used Uniform Manifold Approximation and Projection^100, 101^ (UMAP) to project the transcriptional states for each cell into two-dimensional space (FIG. 3G and Supplementary FIG. S6C). We see a clear separation of the features for the cell line versions in this space, with minimal overlap (FIG. 3G, *right*). Pairwise distances between the centroids of the single-cell features quantify transcriptomic similarities between the cell line versions (FIG. 3H). We see that PC9-VU and PC9-BR1 are more similar to each other than either is to PC9-MGH, which is unsurprising given that PC9-BR1 was derived from PC9-VU (Supplementary FIG. S2B). These results are further supported by alternative dimensionality reduction techniques (Supplementary FIG. S7) and bulk RNA expression data (Supplementary FIG. S8).

The results above illustrate that genetic differences among cell line versions translate to epigenetic differences at the transcriptomic level. However, this genetic-to-epigenetic connection is not one-to-one, i.e., most genetic mutations do not result in an altered expression level of that gene^102^. Instead, mutations that modify protein structure, for example, can alter expression of downstream genes^103^ within the same or related pathways due, e.g., to various feedback mechanisms. Thus, rather than comparing genomic mutations and differential gene expression on a gene-by-gene basis, we establish a baseline for genetic-to-epigenetic connection in terms of biological *process* and molecular *function* within the cell line versions. We use a semantic similarity metric^104^ for comparing gene ontology (GO) terms^105, 106^ for high consequence genetic variants (*low*, *moderate*, and *high* IMPACT score; FIG. 3C) and significantly differentially expressed genes (DEGs; p<0.05) at the transcriptomic level (FIG. 3G). For each cell line version, we calculate pairwise similarity scores between GO terms for each genomic variant and each DEG based on their “distance” within the GO term network^104^ (a directed acyclic graph). Terms within closely related branches of the network are more similar in terms of biological process or function and, hence, receive higher scores than those in different branches (see Methods and Supplementary Note for further details). An aggregate score (between 0 and 1) is then obtained that quantifies the overall relationship between the genetic and transcriptomic states of the cell line versions. For all cell line versions, we get semantic similarity scores close to 0.4 and 0.3 for the “Biological Process” and “Molecular Function” GO categories^105, 106^, respectively (FIG. 3I). These are used as benchmarks against which genetic-to-epigenetic connections for single cell-derived PC9-VU sublines can be compared (see below).

### Single cell-derived sublines are genomically similar but transcriptomically distinct

We perform the same genomic and transcriptomic analyses as above on four PC9-VU sublines (DS3, DS6, DS7, DS9) that exhibit differential responses to EGFRi (FIG. 2C,D). Analysis of the total number of mutations by chromosome (FIG. 4A) shows significantly less variability in the variant count for the sublines relative to the cell line versions (average CV per chromosome = 6.05 vs. 12.84; cf. FIG. 3A). Additionally, unlike the cell line versions, all sublines exhibit similar proportions of unique mutations (FIG. 4B) and numbers of IMPACT mutations (FIG. 4C), with only about 1% of unique mutations predicted to be impactful in all cases. Mutational significance analysis using the same genomic signature as for the cell line versions (see FIG. 3D) shows fewer total and impactful mutations in the sublines (FIG. 4D). This is true for the cancer-associated genes as well (FIG. 4E). The sublines also exhibit a nearly identical mutation class distribution (FIG. 4F). Taken together, these genomic data (FIG. 4A–F) illustrate that there is significantly less genomic variability among the four PC9-VU sublines than among the cell line versions. From the perspective of the G/E/S framework (FIG. 1A), this provides support for the view that the sublines are, in fact, *genetically identical* and represent basins within a common epigenetic landscape defined by the genetic state of PC9-VU.

**Figure 4.**
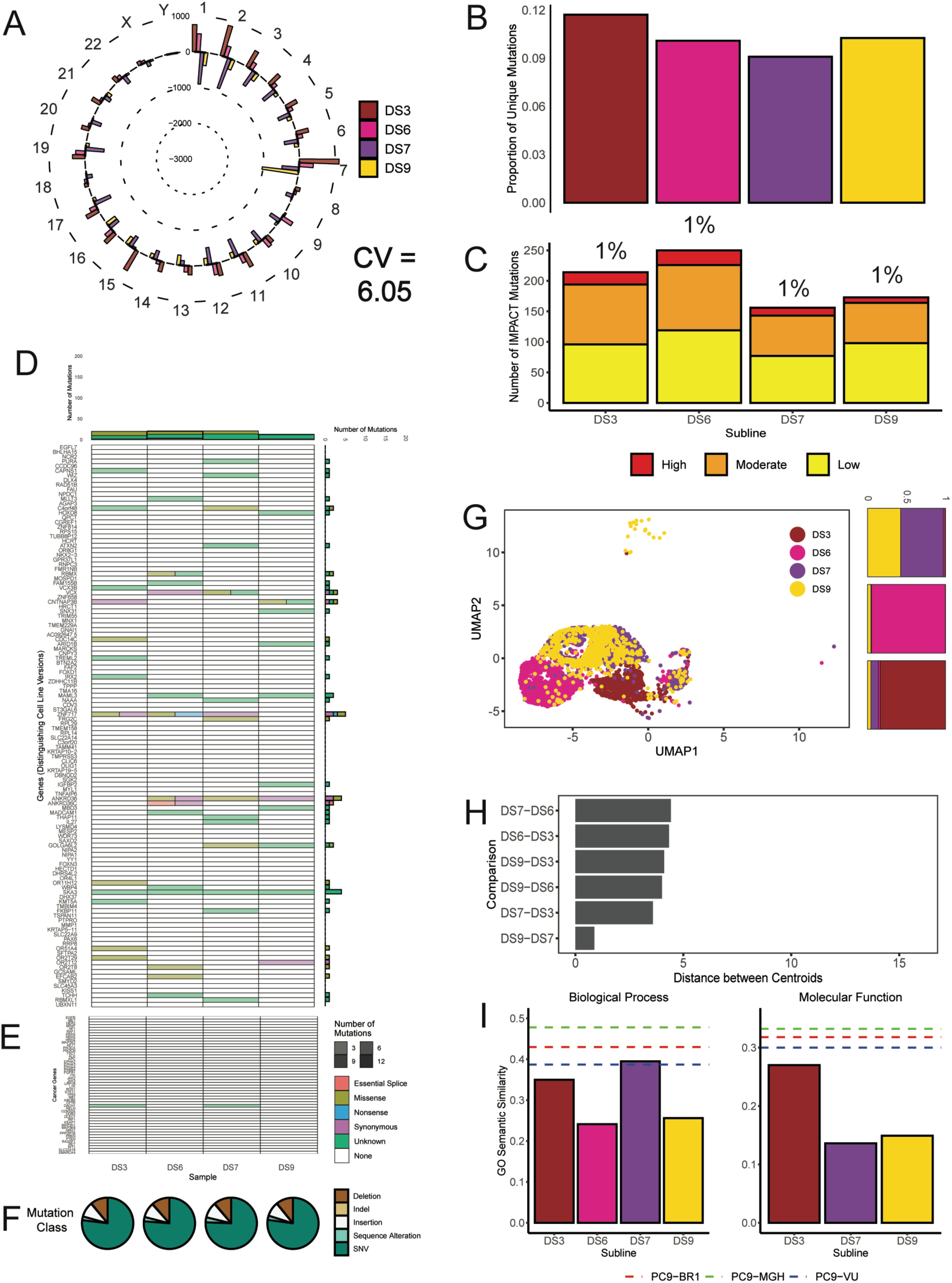
Genomic and transcriptomic characterizations of PC9-VU discrete sublines. (A) Mean-centered mutation count by chromosome for four (of the seven) sublines. Average coefficient of variation across all chromosomes is noted. (B) Proportions of unique mutations in cell sublines. (C) Numbers of IMPACT mutations unique to each subline, stratified by IMPACT classification (*low*, *moderate*, *high*). Percentage of unique IMPACT mutations relative to the total number of unique mutations in each subline is denoted above each bar. (D) Quantification of mutational differences between sublines based on genes that distinguish cell line versions. Heatmap elements are colored based on type of mutation (see FIG. 3D). (E) Same as *D*, but for a literature-curated set of cancer-associated genes implicated in lung cancer. (F) Mutation class pie charts (see FIG. 3F). (G) UMAP visualization of single-cell transcriptomes for sublines plotted in the common UMAP space of all eight PC9 samples (including cell line versions and PC9-VU sublines; see Methods). Proportions of cells in defined clusters (k=3; only points with UMAP1 < 0 and UMAP2 < 5 were considered) are shown as bar plots to the right, colored by subline (*cluster 1*: 0.5% DS3, 2.2% DS6, 54.8% DS7, 42.5% DS9; *cluster 2*: 0.2% DS3, 94.5% DS6, 1.0% DS7, 4.3% DS9; *cluster 3*: 83.6% DS3, 2.4% DS6, 9.7% DS7, 4.3% DS9). (H) Pairwise (Euclidian) distances between the centroids of the transcriptional features in *G* for all sublines. (I) GO semantic similarity analysis of DEGs and unique IMPACT mutations in each subline for “Biological Process” (*left*) and “Molecular Function” (*right*) ontology types. DS6 is not included for “Molecular Function” because it did not produce enough significant GO terms to render a value. Semantic similarity scores for each cell line version (FIG. 3G) are shown as dashed horizontal lines for reference.

Comparing single-cell transcriptomes (FIG. 4G), we see distinctions among the sublines but to a much lesser extent than among the cell line versions. Interestingly, two of the sublines (DS7 and DS9) overlap almost entirely in transcriptional space, despite the fact that they were independently cloned and cultured. A k-means clustering analysis (k=3) shows a near-equivalent distribution of DS7 and DS9 in one cluster, while DS3 and DS6 are highly represented in their own clusters, with some degree of overlap from the other sublines (FIG. 4G, *right*). This is in contrast to the cell line versions, where there is much less overlap among clusters and virtually none between PC9-MGH and PC9-VU/PC9-BR1 (FIG. 3G, *right*). Pairwise distances between subline centroids shows a separation between DS3, DS6, and DS7/DS9 but essentially no separation between DS7 and DS9 (FIG. 4H). Importantly, the distances between subline centroids are all markedly smaller than the distances between centroids for the cell line versions (compare FIGS. 3H and 4H). From the perspective of the G/E/S framework (FIG. 1A), these results suggest that DS7 and DS9 may actually occupy the same basin within the PC9-VU epigenetic landscape while DS3 and DS6 likely occupy different basins. Also note that the subline transcriptomes overlap almost entirely with the parental PC9-VU transcriptome (Supplementary FIG. S7A), as expected, supporting the view that the cell line versions harbor multiple epigenetic states (basins). Moreover, the overlap between the transcriptomic features for the sublines suggests that phenotypic state transitions are occurring between these states. These results are further supported by alternative dimensionality reduction methods (Supplementary FIG. S7B,C) and bulk RNA sequencing (Supplementary FIG. S8).

Semantic similarity analysis between GO terms associated with genetic variants and differentially expressed genes in each of the four sublines (FIG. 4I) reveals a much weaker genetic-to-epigenetic connection as compared to the cell line versions (FIG. 3I). In almost every case (with one exception, DS7: “Biological Process”), the semantic similarities for the sublines are significantly lower than those for all three cell line versions. This can be seen as evidence that the small genomic variations present in the sublines *do not* translate to the transcriptomic level and, therefore, further supports the view from the perspective of the G/E/S framework (FIG. 1A) that the sublines are genetically identical and represent distinct epigenetic states within a common epigenetic landscape.

### Stochastic birth-death simulations suggest sublines are epigenetically monoclonal

The PC9-VU sublines exhibit variability in drug response, not just at the population level (FIG. 2C) but also at the clonal level, as evidenced by variable colony growth in cFP assays (FIG. 5A and Supplementary FIG. S9A) and quantified as distributions of DIP rates (FIG. 2D). To explore the origin of this intra-clonal variability, we perform stochastic simulations^107^ on a simple birth-death model of cell proliferation to ascertain whether intrinsic noise alone is sufficient to explain the experimental observations (see Methods). We perform a battery of *in silico* cFP assays where untreated single cells grow into colonies of variable size at a fixed proliferation rate (division rate – death rate) and are then treated with drug, modeled by reducing the proliferation rate. Colony sizes are tracked over time (FIG. 5B and Supplementary FIG. S9B) and DIP rate distributions are calculated and statistically compared against experimental distributions (FIG. 5C and Supplementary FIG. S9C). We repeat this procedure for a wide range of division and death rates to identify ranges of parameter values that can statistically reproduce (p>0.1) experimental DIP rate distributions. For all PC9-VU sublines (except DS8, see below), we find ranges of parameter values that are physiologically plausible (FIG. 5D and Supplementary FIG. S9D). The fact that a birth-death model with fixed rates of division and death, when simulated stochastically, can reproduce experimental DIP rate distributions supports the view that these sublines (DS1, DS3, DS4, DS6, DS7, DS9) are, in fact, epigenetic clones, i.e., individual basins within the PC9-VU epigenetic landscape (FIG. 1A). Moreover, that many values of division and death rates can reproduce experimental results is consistent with the idea that the molecular states of cells fluctuate within basins of attraction (FIG. 1A). In other words, rates of division and death themselves fluctuate because they are fundamentally driven by biochemical signaling pathways that are susceptible to stochastic fluctuations in intracellular species concentrations^20, 21, 23, 85–87^.

**Figure 5.**
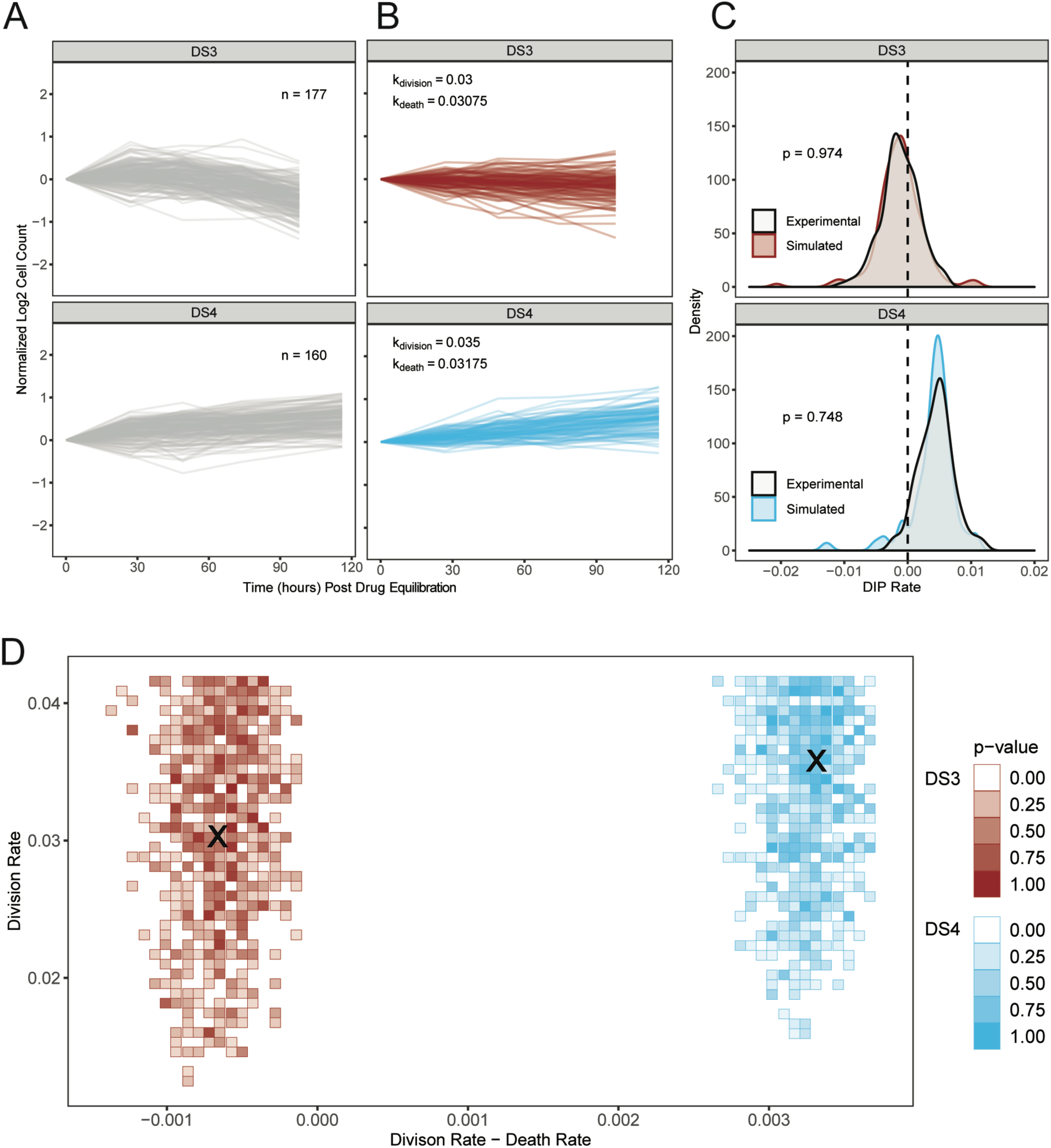
Stochastic simulations of a simple birth-death model reproduce DIP rate distributions from PC9-VU sublines. (A) Experimental cFP time courses for two representative sublines (DS3 and DS4) in response to 3μM erlotinib (same data used to generate the corresponding DIP rate distributions in FIG. 2D). Each trace corresponds to a single colony, normalized to 72h post-drug treatment. Only colonies with initial cell counts greater than 50 at the time of treatment are shown. (B) *In silico* cFP time courses with division and death rate constants that closely reproduce the experimental time courses in *A*. Trajectories are normalized to the time at which the simulated drug treatment was initiated, simulated cell counts are plotted only at experimental time points, and only colonies with initial cell counts greater than 50 at the time of simulated drug treatment are shown. (C) Comparison of experimental and simulated DIP rate distributions from calculated from time courses in *A* and *B*. Distributions are compared statistically using the Kolmogorov-Smirnov (K-S) test^153^. Dashed black line signifies zero DIP rate, for visual orientation. (C) Parameter scan of division and death rate constants for the two sublines in *A*–*C*. For each pair of rate constants, we ran the same number of model simulations as the associated experimental cFP time courses in *A*, calculated DIP rates and compiled them into a distribution, and then statistically compared against the corresponding experimental DIP rate distribution using the K-S test. Color shades correspond to p-values; all p<0.1 are colored white, indicating lack of statistical correspondence to experiment. **×** denotes a division and death rate constant used in *B*.

### One single cell-derived subline likely harbors a second genetic state

As mentioned above, the PC9-VU subline DS8 differs significantly from the other sublines, exhibiting near total resistance to EGFRi (FIG. 2C) and a bimodal DIP rate distribution under cFP (FIG. 2D). To investigate the origin of these differences, we perform on DS8 the same multi-omic and simulation analyses as reported above (FIGS. 3–5). On a per chromosome basis, DS8 tends to have more mutations than the average subline (although this is true of DS3 and DS6 as well; Supplementary FIG. 10A). It also has a larger proportion of unique mutations relative to all other sublines (FIG. 6A) and substantially more unique IMPACT mutations (total and as a percentage; FIG. 6B). In terms of mutational significance and class distribution (Supplementary FIG. S10B–D), DS8 does not differ significantly from the other sublines. However, it is important to note that DS8 does *not* have the same resistance conferring mutation (EGFR-T790M) as PC9-BR1 (Supplementary FIG. S10C), indicating a different (unknown) resistance mechanism is at play (see Supplementary Note for further discussion). Transcriptomically, DS8 differs significantly from the other sublines (FIG. 6C,D) and cell line versions (Supplementary FIG. S7A). This is evident in other dimensionality reduction projections as well (Supplementary FIG. S7B,C) and bulk RNA expression profiles (Supplementary FIG. S8). GO semantic similarity analysis shows a strong genetic-to-epigenetic connection in DS8 (FIG. 6E), on par with the baselines set by the cell line versions (FIG. 3I) and significantly higher than the other sublines (FIG. 4I).

**Figure 6.**
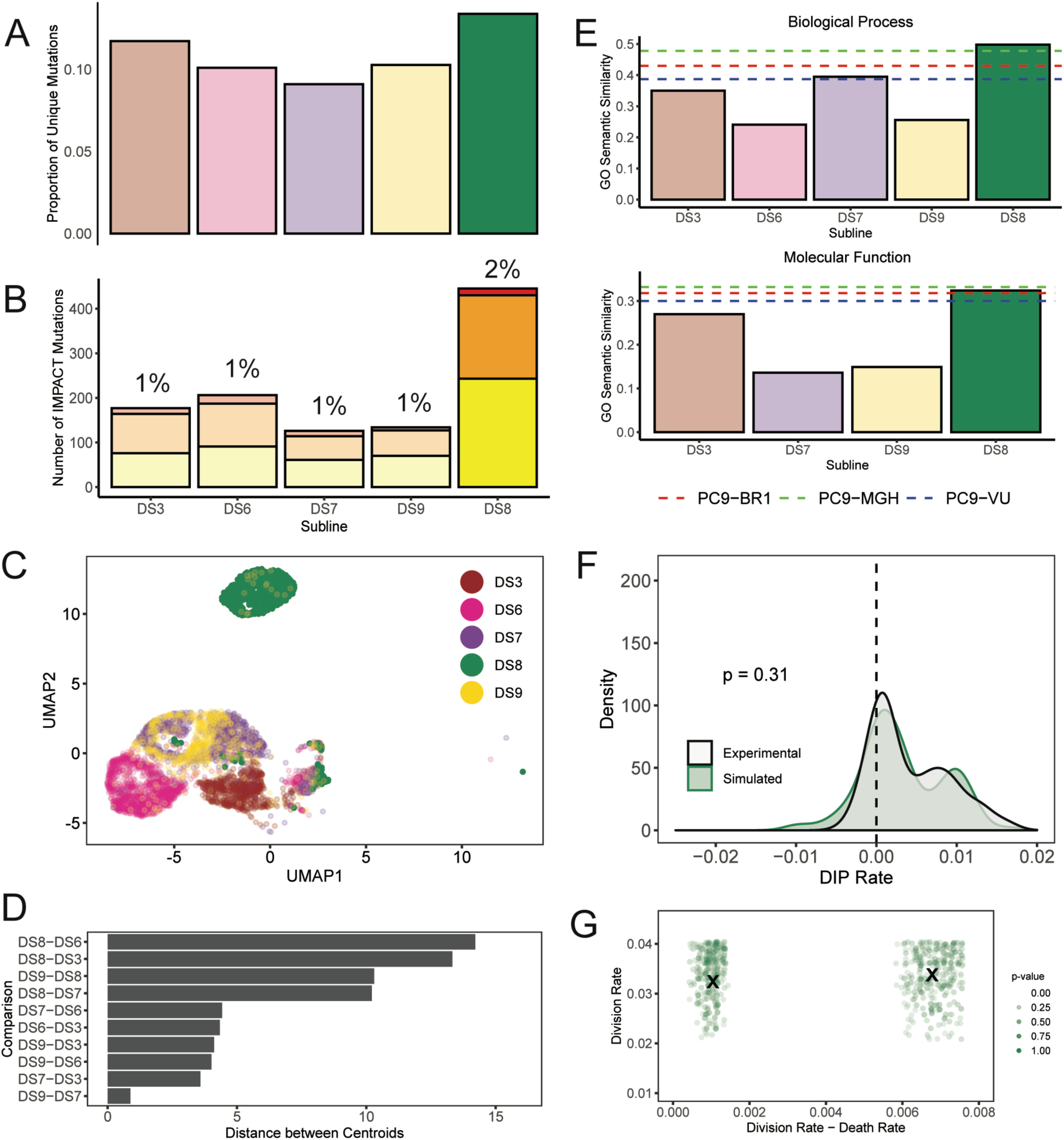
Genomic and transcriptomic analyses of DS8 identify a genetically-distinct state. (A) Proportions of unique mutations in all PC9-VU sublines, including DS8. Bars for all other sublines are displayed with reduced transparency to emphasize the distinction with DS8. (B) Numbers of IMPACT mutations unique to each subline, stratified by IMPACT classification (*low*, *moderate*, *high*). Percentage of unique IMPACT mutations relative to the total number of unique mutations in each subline is denoted above each bar. Again, DS8 is emphasized. (C) UMAP visualization of single-cell transcriptomes for all sublines, including DS8. Sublines are plotted in the common UMAP space of all samples, with DS8 highlighted. (D) Pairwise (Euclidian) distances between the centroids of the transcriptional features in *C*, including distances from DS8. (E) GO semantic similarity analysis of sublines, including DS8, for “Biological Process” (*top*) and “Molecular Function” (*bottom*) ontology types. DS6 is not included for “Molecular Function” because it did not produce enough significant GO terms to render a value. The semantic similarity scores for each PC9 cell line version (FIG. 3I) are shown as dotted horizontal lines for reference. DS8 is highlighted for emphasis. (F) Comparison of experimental and simulated DIP rate distributions for DS8. Statistical analysis was performed using a K-S test. Dashed line signifies zero DIP rate for visual orientation. (G) Parameter scan of division and death rate constants for a two-state birth-death model. A four-dimensional (two division-death rate constant pairs) parameter scan was performed and projected into two dimensions. **×** denotes parameter values used to generate the simulated DIP rate distribution in *F*.

Taken together, these multi-omics data (FIG. 6A–E and Supplementary FIGS. S7, S8, and S10A–D) strongly suggest that DS8 is *genetically distinct* from the other PC9-VU sublines and from the other cell line versions (PC9-MGH and PC9-BR1). We further support this claim by performing stochastic simulations on an expanded version of the birth-death model described above that contains two clones with distinct rates of division and death (see Methods and Supplementary Note). As for the other sublines, we perform *in silico* cFP assays (Supplementary FIG. S10E) and compare the simulated DIP rate distributions against the bimodal distribution seen experimentally (FIG. 6F). We find ranges of rate parameter values that can statistically reproduce the experimental result (FIG. 6G). Note that the mathematical model is agnostic as to whether or not the two *in silico* clones are genetically distinct (i.e., it simply posits that the clones have distinct proliferation rates). Thus, while the weight of the evidence points to DS8 as harboring a distinct genetic state, it is an open question as to whether that state emerged prior to establishment of the subline or subsequently (Supplementary FIG. S11; see below for further discussion).

## DISCUSSION

Many modern cancer therapies have focused on targeting specific genetic mutations within a tumor. However, recent studies have shown that a complex interplay between genetic and non-genetic factors likely plays a key role in targeted treatment failure^8, 96^. Genetic and non-genetic heterogeneities in cancer have been extensively studied over the past decade or more^5, 20, 22, 52, 108–111^, but most studies have lacked a clear theoretical framework to understand them jointly. Here, we have utilized the framework of Waddington^34^, extended by Huang^13, 15, 19, 35, 36, 64^, Polyak^8, 16, 17^, and others^41, 48^, to deconvolve the roles of genetic, epigenetic, and stochastic variabilities on drug response phenotypes. This G/E/S framework (FIG. 1) posits that tumors are composed of multiple genetic states, each of which emanates an epigenetic landscape comprising multiple phenotypes (basins of attraction), within which cell states fluctuate due to intrinsic and extrinsic sources of noise that can occasionally drive phenotypic state changes.

To provide substantive experimental support for this framework, we used an *in vitro* tumor heterogeneity model comprising multiple versions of the NSCLC cell line PC9 (VU, MGH, BR1) that were propagated separately from each other over extended periods of time (Supplementary FIG. S2B) and exhibit drastically different drug responses (FIG. 2A,B). We made the forthright assumption that the cell line versions are genetically distinct (Supplementary FIG. S1) and used whole-exome sequencing to establish a baseline for mutational differences, which were significant (FIG. 3A–F). According to the G/E/S framework, the PC9 cell line versions lie along a “genetic axis” (FIG. 7A, *left*), with PC9-BR1 having been derived from PC9-VU (FIG. 7B), and each has an associated epigenetic landscape comprising multiple discrete phenotypic states (FIG. 7A, *right*). These epigenetic landscapes are revealed as distinct transcriptional features by sc-RNA-seq (FIG. 3G,H) and their connections to the underlying genetic states are established by semantic similarity analysis between mutated and differentially-expressed genes (FIG. 3I). However, individual basins are not discernable in the parental PC9 transcriptional features (FIG. 3G). According to the theory, this can be explained by cell-state fluctuations that “smear” the individual features together (FIG. 1A, *right*). This view is substantiated by sc-RNA-seq for four single cell-derived PC9-VU sublines whose transcriptomic features overlap significantly with different portions of the parental PC9-VU feature (FIG. 4G,H and Supplementary FIG. S7A). Moreover, overlap among the subline transcriptomic features provides some evidence for fluctuation-driven phenotypic state changes. Genomic differences among these sublines (FIG. 4A–F) are substantially smaller than the baseline established by the cell line versions, as is the genetic-to-epigenetic connection (FIG. 4I), further supporting the assertion that these sublines are genetically identical (Supplementary Table S1) and constitute basins of a common epigenetic landscape (FIG. 7A, *right*). Differential drug responses across the PC9-VU sublines (FIG. 2C) can be attributed to the different molecular (transcriptomic) states of the sublines (FIG. 4G; except for DS8, see below) and drug-response variability seen at the clonal level within the sublines (FIG. 2D) can be similarly attributed to fluctuations in molecular states, captured using stochastic simulations at the cell population level (FIG. 5 and Supplementary FIG. S9).

**Figure 7.**
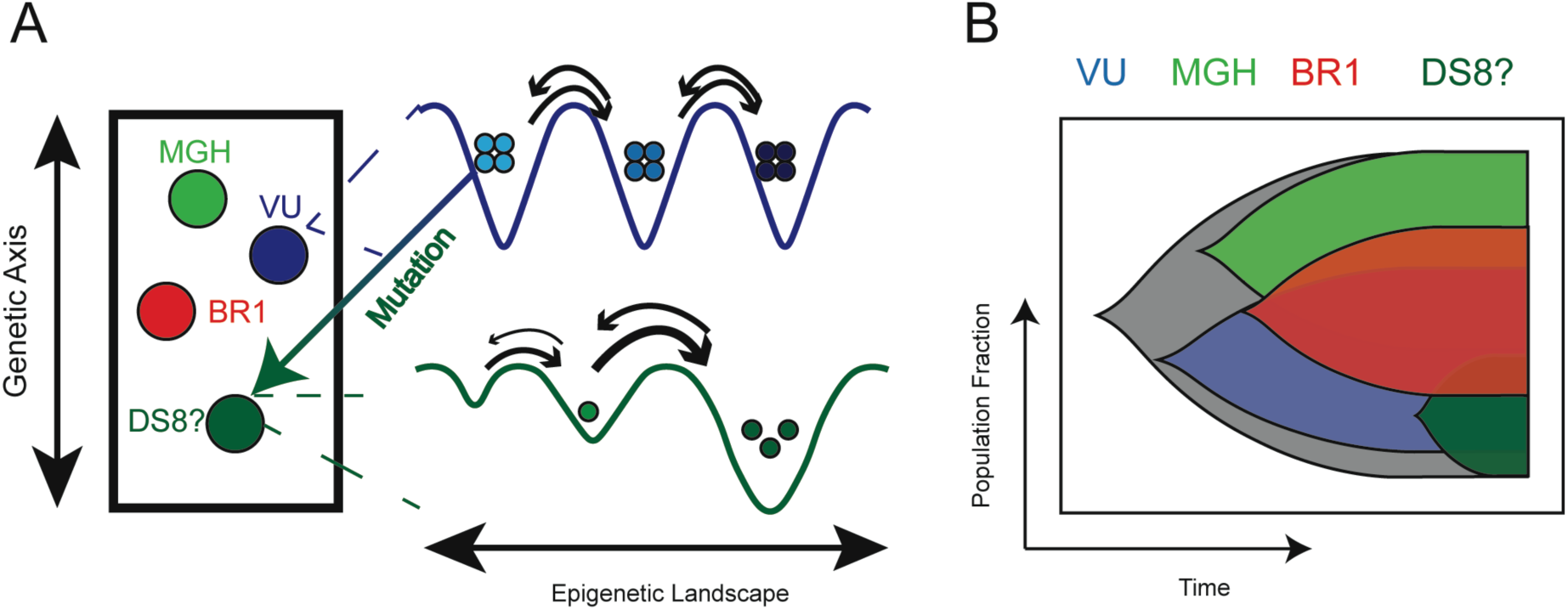
Application of the G/E/S heterogeneity framework to a hypothetical tumor composed of PC9 cell line family members. (A) Cell line versions PC9-MGH (*light green*), PC9-BR1 (*red*), and PC9-VU (*blue*) fall in different regions of the genetic axis (*left*). Each of these versions has a corresponding epigenetic landscape (only PC9-VU shown; *right*). Due to a mutation in PC9-VU (denoted by the arrow) a new genetic state emerged (dark green), which we hypothesize is the origin of DS8. The new genetic state also has an associated epigenetic landscape with basins that appear as a large shoulder in the positive region of the bimodal DIP rate distribution for DS8 (FIGS. 2D and 6F). It is still unclear whether DS8 comprises one or multiple genetic states (see Supplementary FIG. S11 and Supplementary Note). (B) Schematic view of genetic clonal evolution for the PC9 cell line family. From the recent common ancestor (grey; assumed to be from the original PC9 patient tumor), PC9-VU and PC9-MGH evolved independently in the absence of selective pressures. PC9-BR1 was derived from PC9-VU and acquired mutations (including, but not limited to EGFR-T790M) in the presence of selective pressure (i.e., dose-escalation therapy). The DS8 subline contains a distinct genetic state that emerged from PC9-VU in the absence of selective pressure and is resistant to EGFRi (absent EGFR-T790M).

Throughout our analyses, DS8 was a persistent outlier among the PC9-VU sublines: it is almost entirely resistant to EGFRi (FIG. 2C and Supplementary FIG. S4B); it displays a bimodal DIP rate distribution at the clonal level (FIG. 2D and FIG. 6F,G); it has significantly more unique and IMPACT mutations than the other sublines (FIG. 6A,B); its transcriptomic state is substantially distinct from the other sublines (FIG. 6C,D), on par with the other cell line versions (Supplementary FIG. S7A); and its genetic-to-epigenetic similarity scores are comparable to those of the cell line versions (FIG. 6E). Furthermore, DS8 does not carry the classic EGFR-T790M mutation that confers EGFRi resistance, as found in PC9-BR1 (Supplementary FIG. S10C). All this points to DS8 harboring a distinct genetic state that emerged out of PC9-VU at some point in the past in the absence of selective pressure (FIG. 7B). Within the G/E/S framework, this places DS8 on the genetic axis alongside the other cell line versions (FIG. 7A, *left*) and asserts the existence of an associated epigenetic landscape (FIG. 7A, *right*). An open question remains, however, as to whether the DS8 genetic state emerged subsequent to establishment of the subline, in which case DS8 would likely comprise *two* distinct genetic states (the mutant and the PC9-VU state from which it arose; Supplementary FIG. S11A), or prior to its establishment, which would stipulate that mutant cells also remain in the parental line (Supplementary FIG. S11B). The bimodal DIP rate distribution for DS8 seems to favor the former scenario: the left mode of the distribution largely overlaps with the PC9-VU distribution, while the right mode, presumably associated with the new genetic state, is distinct, lying between PC9-VU and PC9-BR1 (FIG. 2B and Supplementary FIG. S4C). However, we cannot entirely discount the latter possibility since a small proportion of PC9-VU and DS9 cells reside in the same region of transcriptional space as DS8 (FIGS. 3G, 4G, and 6C). It is possible, therefore, that the emergent DS8 genetic state associates with an epigenetic landscape comprising basins lying in distant regions of transcriptional space (Supplementary FIG. S11B; see Supplementary Note for further discussion). Whatever the case, the weight of the evidence reported here supports the view that DS8 is genetically distinct from the other PC9 cell line versions and sublines and has access to phenotypic states that are either completely or largely unavailable to PC9-VU, consistent with the G/E/S perspective on the complex interplay between genetic and non-genetic heterogeneities in cancer (FIG. 1).

In addition to the experimental evidence presented here, the G/E/S framework is wholly consistent with numerous prior reports in the literature. Ben-David et al.^42^ showed that numerous “strains” (comparable to our cell line versions) of human cancer cell lines, obtained from different institutions, display extensive genetic heterogeneity and that genetically similar strains exhibit similar transcriptomic signatures and drug-response profiles. Thus, they argued that cancer cell lines can drift genetically when kept in culture independently, consistent with our results for the PC9 cell line versions (FIG. 3). Moreover, the spontaneous emergence of the drug-resistant DS8 genetic state out of PC9-VU in the absence of selective pressure (FIG. 6) aligns with observations by Ramirez et al.^2^ and Hata et al.^28^, who independently reported diverse resistance-conferring mutations arising in drug-treated and untreated PC9-MGH clones. Shaffer et al.^112^ described a transient, transcriptionally-encoded pre-resistance state in two BRAF-mutant melanoma cell lines that they attributed to epigenetic mechanisms (both gene expression states and chromatin accessibility). Although presented phenomenologically, we can interpret their results through the lens of the G/E/S framework: similar to our single cell-derived sublines, the observed pre-resistant state can be viewed as a basin within a BRAF-mutant melanoma epigenetic landscape that, based on its molecular make up, is insensitive to BRAF inhibition. Moreover, the observation that individual cells transition into and out of this state is consistent with the idea that stochastic fluctuations drive phenotypic state transitions over epigenetic landscapes. Since only a small minority of cells seem to have access to the pre-resistant state at any given time^112^, we can infer that the basin is shallow because outward transitions would be easily driven by intracellular noise. This view is consistent with seminal works by Arkin and co-workers^21, 23, 75, 85, 86, 113^, who detailed the role of gene expression noise in probabilistic cell fate decisions and variations in phenotypic identity using stochastic simulations and experimentation. Elowitz et al.^52^ demonstrated how noise in transcription and translation can lead to phenotypic switching in bacterial populations. Similar behaviors have been observed by Raj and van Oudenaarden in yeast^20^ and more recently by Magklara and Lomvardas^71^ in mammalian systems. In cancer, cell-to-cell variability in protein concentrations^114^ and stochastic phenotypic state switching^59, 115^ have been proposed as survival mechanisms against treatment, in line with our observations of unimodal DIP rate distributions and small proportions of positive-DIP rate clones in isogenic PC9 cell populations (FIGS. 2 and 5 and Supplementary FIGS. S3, S4, and S9).

It is now abundantly clear that therapies based on evaluating tumor genetics alone cannot solve the growing problems of increasing rates of cancer incidence and mortality worldwide^116, 117^. The G/E/S framework presented here offers an alternative to the traditional gene-centric view of cancer that may transform how we understand and treat the disease. For example, that each genetic state emanates an epigenetic landscape with numerous accessible phenotypic states may explain why genetic screening alone has not produced effective, durable anticancer therapies: some phenotypes may have molecular compositions that enable their survival under drug treatment. Cells pre-existing in these states (e.g., that observed by Shaffer et al.^112^), and/or those that escape into them upon drug addition, may act as a refuge from which genetic resistance mutations can arise. This “bet hedging” strategy of non-genetic phenotypic diversification in the absence of stressors has been observed in bacteria^109, 113, 118, 119^ and proposed as a survival strategy in cancer^30, 62^. Furthermore, alternative treatment strategies have been proposed that focus on identifying and targeting so-called “cancer stem cells”^120^ (CSCs), rare cell types that do not differ genetically from bulk tumor cells but are phenotypically distinct and capable of seeding new tumors. Unfortunately, these treatments have so far been unsuccessful^121^. The G/E/S framework provides a possible explanation as to why: CSCs can be viewed as residing in a shallow basin within an epigenetic landscape; killing cells in this basin does not eradicate the basin, leaving it available to be repopulated by cells from “adjacent” basins. This suggests that a better approach is not to target individual phenotypic states but rather to target the epigenetic landscape as a whole^122^. This process, termed “targeted landscaping”^30, 43^, would aim to use drugs, in combination or succession, to alter the topography of a landscape to favor transitions out of drug-tolerant states and into drug-sensitive states. The rationality of this approach is supported by multiple studies, including in cancer, where resistance to one drug has been shown to confer sensitivity to a different drug (or drug class), commonly referred to as “collateral sensitivity”^123–125^. Several studies have also demonstrated that alternative drug scheduling regimens may have benefits compared to traditional single-agent applications or up-front combinations^45, 126^. Our work suggests extending these approaches to account for all levels of heterogeneity in tumors, i.e., genetic profiling to enumerate dominant genetic states, followed by characterization of the associated epigenetic landscapes using single-cell experimental approaches, possibly aided by advanced theoretical and computational modeling techniques^41, 64, 65, 73, 127–132^. These landscapes can then be screened against batteries of available drugs, *in vitro* and *in silico*, to devise individualized treatments that can be tested against patient-derived xenografts *in vivo* before being administered clinically. This approach would leverage novel technologies and currently available drugs to offer the possibility of profound improvements to anticancer therapies.

## METHODS

### Cell culture and reagents

PC9-VU and PC9-BR1 were obtained as a gift from Dr. William Pao (Roche, formerly of Vanderbilt University Medical Center). PC9-MGH was obtained as a gift from Dr. Jeffrey Settleman (Massachusetts General Hospital). PC9-VU, PC9-MGH, and PC9-BR1 were individually fluorescently labeled with histone H2B conjugated to monomeric red fluorescent protein (H2BmRFP), as previously described^30, 81, 88, 89, 133^. The PC9 cell line versions and derivatives were cultured in Roswell Park Memorial Institute (RPMI) 1640 Medium (Corning) with 10% Fetal Bovine Serum (Gibco). Cells were incubated at 37°C, 5% CO_2_, and passaged twice a week using TrpLE (Gibco). Cell lines and sublines were tested for mycoplasma contamination using the MycoAlert^TM^ mycoplasma detection kit (Lonza), according to manufacturer’s instructions, and confirmed to be mycoplasma-free. Cell line identity was confirmed using mutational signatures in whole exome sequencing. Approximately 99% of the PC9-VU mutations were also seen in PC9-MGH, but PC9-MGH exhibited a large number of unique and total mutations independently (Supplementary FIG. S5C). Additionally, using the *SNPRelate* R package (version 1.18.1), sample genotypes were projected into low dimensional space and clustered based on similarity (Supplementary FIG. S12). PC9-VU and PC9-MGH clustered together and were represented closely in reduced dimensional space, further suggesting that they arose from the same parent cell population. Erlotinib was obtained from Selleck Chemicals (Houston, TX) and solubilized in dimethyl sulfoxide (DMSO) at a stock concentration of 10mM and stored at −20°C. Cell lines were originally stored at −80°C, then moved into liquid nitrogen.

### Derivation of single-cell sublines

The PC9-VU sublines were generated by limiting dilution of the parental cell line in 96-well plates. Wells with single cells were expanded for multiple weeks until large enough to be saved as frozen cell stocks. A single stock of each subline was brought back into culture, passaged for two weeks, and used for drug-response experiments. After 3-4 weeks of continued passaging, sublines were used for whole exome sequencing, bulk RNA sequencing, and single-cell RNA sequencing experiments. Since sublines were isolated from PC9-VU, they retained the same H2BmRFP nuclear label as cell line versions.

### Population-level DIP rate assay

Cells were seeded in black, clear-bottom 96-well plates (Falcon) at a density of 2500 cells per well with six replicates for each sample. Plates were incubated at 37°C and 5% CO_2_. After cell seeding, drug was added the following morning and was changed every three days until the end of the experiment or confluency. Untreated samples were allowed to grow in DMSO-containing media until confluency, with media changes every three days. Plates were imaged using automated, fluorescence microscopy (Cellavista Instrument, Synentec). Twenty-five non-overlapping fluorescent images (20X objective, 5×5 montage) were taken twice daily for a total of 500 hours, or until confluency. Cellavista image segmentation software (Synentec) was utilized to calculate nuclear count (i.e., cell count) per well at each time point (Source = Cy3, Dichro = Cy3, Filter = Texas Red, Emission Time = 800μs, Gain = 20x, Quality = High, Binning = 2×2). Cell nucleus count across wells was used to calculate mean and 95% confidence intervals and normalized to time of drug treatment. Data was visualized using the *ggplot2* R package (version 3.2.0).

### Clonal Fractional Proliferation assay

Cells were single-cell seeded in a black, clear-bottom 384-well plate (Greiner) using fluorescence-activated cell sorting (FACS Aria III, RFP^+^). Plates were incubated at 37°C, 5% CO_2_, and cells were allowed to grow into small colonies over eight days in complete media (no media change). Drug was then added to cell colonies and changed every three days. Plates were imaged using the Cellavista Instrument (Synentec). Nine non-overlapping fluorescent images (3×3 montage of the whole well at 10X magnification) were taken once daily for a total of seven days. Cellavista image segmentation software (Synentec) was utilized to calculate nuclear count (i.e., cell count) per well at each time point (Source = Cy3, Dichro = Cy3, Filter = Texas Red, Emission Time = 800μs, Gain = 20x, Quality = High, Binning = 2×2). Depending on the number of wells that passed quality control thresholding (at least 50 cells per colony at the time of treatment), 160-280 replicates were included for each sample (FIG. 5A and Supplementary FIGS. S3B,D,E, S4A, S9A, and S10E). DIP rates were calculated from 48h post-treatment to the end of the experiment (single-colony traces in FIG. 5A and Supplementary FIGS. S3D-E, S4A, S9A, and S10E; linear model fits in Supplementary FIG. S3B) using the *lm* function in R. DIP rates for each sample were combined and plotted as a kernel density estimate (FIGS. 2D, 5C, and 6F and Supplementary FIGS. S4C and S9C). Mood’s median test^134^ was performed to determine statistical difference between subline DIP rate distributions (Supplementary FIG. 4C) using the *RVAideMemoire* R package. Data was visualized using the *ggplot2* package.

### DNA and RNA extraction

Genomic DNA was extracted using the DNeasy Blood and Tissue Kit (Qiagen), according to the manufacturer’s protocol. Total RNA was extracted using a Trizole extraction (ThermoFisher), according to the manufacturer’s protocol.

### DNA bulk whole exome sequencing

Libraries were prepared using 150 ng of genomic DNA by first shearing the samples to a target insert size of 200 bp. Illumina’s TruSeq Exome kit (Illumina, Cat: 20020615) was utilized per manufacturer’s instructions. The samples were then captured using the IDT xGen® Exome Research Panel v1.0 (IDT, Cat: 1056115). The resulting libraries were quantified using a Qubit fluorometer (ThermoFisher), Bioanalyzer 2100 (Agilent) for library profile assessment, and qPCR (Kapa Biosciences Cat: KK4622) to validate ligated material, according to the manufacturer’s instructions. The libraries were sequenced using the NovaSeq 6000 with 150 bp paired end reads. RTA (version 2.4.11; Illumina) was used for base calling and sequence-specific quality control (QC) analysis was completed using MultiQC v1.7. Reads were aligned to the University of California, Santa Cruz (UCSC) hg38 reference genome using ‘BWA-MEM’^135^ (version 0.7.17) with default parameters.

### DNA whole exome sequencing analyses

#### Genomic mutational analysis

Mutation analysis for single nucleotide polymorphisms (SNPs) and insertion/deletions (InDels) was performed using an in-house variant calling pipeline based on the Genome Analysis Toolkit (GATK, Broad Institute) recommendations. Duplicate reads were marked and replaced using PICARD (Broad Institute). Base recalibration and variant calling were performed using GATK version 3.8 (Broad Institute). Variants were selected and filtered based on gold standard SNPs and InDels, as well as a hard filtration according to GATK recommendations (SNPs: QD<2, QUAL<30, SOR>3, FS>60, MQ<40, MQRankSum<-12.5, ReadPosRankSum<-8; InDels: QD<2, QUAL<30, FS>200, ReadPosRankSum<-20). Total variants were counted using VCFtools^136^ (version 0.1.15; Supplementary FIG. S5A). Sequencing metrics were calculated using vcfR^137^ (version 1.8.0). These metrics included read depth, mapping quality, and a Phred quality score^138, 139^ (Supplementary FIG. S5B). Variants were annotated by Variant Effect Predictor^98^ (VEP; Ensembl Genome Browser version 95, available at ensembl.org) with multiple indicators, including chromosome name, gene symbol, mutation class, mutation type, and IMPACT rating (see Methods and Supplementary Note). Variant distribution was organized as a mean-centered mutation count per chromosome for samples within each comparison set (cell line versions – FIG. 3A, sublines – FIG. 4A, sublines with DS8 – Supplementary FIG. S10A). Variants were categorized by mutation class (FIGS. 3F and 4F and Supplementary FIG. S10D). Variant overlap analysis was conducted using VCFtools and visualized using the *UpSet* (version 1.4.0) R package (Supplementary FIG. S5C–E). Variants unique to each sample were plotted as proportions of the total mutations in that sample (FIGS. 3B, 4B, and 6A). One of the VEP annotations used was an IMPACT rating, a variant consequence classifier. Variants were categorized into *modifier* (no evidence of impact), *low* (unlikely to have disruptive impact), *moderate* (non-disruptive but might have effect), and *high* (assumed to have disruptive impact). Impact ratings were identified for mutations unique to each population in the cell line version (FIG. 3B) and subline (FIG. 4B and 6A) comparisons. *Modifier* variants were not plotted but represented a majority of variants in all samples. Variants categorized into *low*, *moderate*, or *high* are hereafter referred to as IMPACT mutations.

#### Automatic generation of a genetic mutational signature

Unique cell line version mutations were input into *dNdScv*^140^ to generate a mutational signature that could define the genetic heterogeneity within the PC9 cell line family. In this analysis, genes are first annotated by type (synonymous, nonsynonymous - missense, nonsense, splice site, unknown; Supplementary FIG. S1). *dNdScv* then uses a maximum-likelihood model to detect genes under positive selection (i.e., potential driver mutations). For each gene, a variety of models are utilized to identify genes that have substantially more nonsynonymous mutations than expected in each of the nonsynonymous mutation types, as compared to synonymous mutational load. These metrics are combined together to calculate a global p-value (see original publication^140^ for details). We use this approach, without the default covariates (because they were not available for hg38) and a maximum number of mutations per gene per sample = 50 (to limit a hyper-mutator phenotype) to determine genes with a global p-value less than 0.05 in the cell line versions. Since we assume these cell lines are genetically distinct, this analysis sets the baseline for the genetic heterogeneity within other cell populations. We visualized the mutation data as a heatmap of these significant genes and cell line versions (FIG. 3D), sublines (FIG. 4D), and sublines with DS8 (Supplementary FIG. S10B), colored by annotated mutation type and number of mutations.

#### Literature-curated, cancer-associated mutational signature

A cancer-associated gene list was established to supplement the predicted genetic heterogeneity signature above. The gene list was created from the NIH Genetics Home Reference (GHR) key lung cancer genes (ghr.nlm.nih.gov/condition/lung-cancer#genes) and two additional publications of key mutations in lung cancer^2, 99^. Associated heatmaps were generated of this gene list for cell line versions (FIG. 3E), sublines (FIG. 4E), and sublines with DS8 (Supplementary FIG. S10C), colored by annotated mutation type and number of mutations.

### RNA single-cell transcriptome sequencing

sc-RNA-seq libraries were prepared using the 10X Genomics gene expression kit (version 2, 3’ counting^141^; Supplementary FIG. S6) and cell hashing^142^ (Supplementary FIG. S13). Cells were prepared according to recommendations from the cell hashing protocol on the CITE-seq website (cite-seq.com/protocol). After cell preparation, 1 ng of eight different cell hashing antibodies (TotalSeq-A025(1-8) anti-human Hashtag, Biolegend) were used to label each of the eight samples (labeling efficiency in Supplementary FIG. S13). Hashed single-cell samples were combined in approximately similar proportions and ‘super-loaded’ (aiming for ∼20,000 cells; ∼15,400 cells were obtained) onto the Chromium instrument. Cells were encapsulated according to manufacturer guidelines. Single-cell mRNA expression libraries were prepared according to manufacturer instructions. The leftover eluent of the mRNA expression library, containing the hashtag oligonucleotides (HTOs), was utilized to further size select the HTO library. The size-selected HTO library was PCR amplified to obtain higher quality reads. Libraries were cleaned using SPRI beads (Beckman Coulter) and quantified using a Bioanalyzer 2100 (Agilent). The libraries were sequenced using the NovaSeq 6000 with 150 bp paired-end reads targeting 50M reads per hashed sample for the mRNA library and a spike-in fraction for the HTO library. RTA (version 2.4.11; Illumina) was used for base calling and MultiQC (version 1.7) for quality control. Gene counting, including alignment, filtering, barcode counting, and unique molecular identified (UMI) counting was performed using the *count* function in the 10X Genomics software *Cell Ranger* (version 3.0.2) with the GRCh38 (hg38) reference transcriptome (Supplementary FIG. S6A). We utilized *CITE-seq-Count* (github.com/Hoohm/CITE-seq-Count) to count hashtags from the HTO library. We then used the *Demux* function in the R package Seurat^143^ (satijalab.org/seurat) to demultiplex the HTO and mRNA libraries, and pair cells to their associated hashtag. Data was integrated into a count matrix with genes and cells, with hashtag identity as a metadata tag.

### RNA single-cell transcriptome analyses

After creating the demultiplexed, single-cell gene expression matrix, we removed multiplets (at least two different hashtags detected with a single cell barcode) from the dataset to leave just singlets or 2+ of the same hashtag. Feature selection was performed (0.1 < average gene expression < 8, log variance-to-mean ratio > 1; 574 genes met criteria; Supplementary FIG. S6B). A cell cycle score was regressed out of the expression matrix, and did not have a large effect on the analysis, at least in terms of sample groupings (Supplementary FIGS. S6C and S7A). Therefore, non-regressed data was used for all visualizations and analyses reported here. Data were visualized using the Uniform Manifold Approximation and Projection (UMAP) dimensionality reduction algorithm (FIGS. 3G, 4G and 6C and Supplementary FIG. S7A) as implemented in the Seurat^143^ R package. To facilitate comparisons across cell line versions and sublines, we performed the UMAP projections in the space of all eight cell line versions and PC9-VU sublines. To quantify overlap of cells between transcriptomic features, we performed k-means clustering (k=3) of the cell line versions (FIG. 3G) and sublines (FIG. 4G) using the *kmeans* function of the R *stats* package (for the sublines, only values with UMAP1 < 0 and UMAP2 < 5 were considered). We also calculated distances between the medoid (centroid) of each sample in a common UMAP space (FIGS. 3H, 4H, and 6D) using the *dist* function (Euclidian) in base R. Differential expression analysis was then performed between a single sample (i.e., PC9-VU) and the rest of the members of the comparison set (i.e., cell line versions, sublines, or sublines including DS8) using the Wilcoxon rank sum test^144^ (see Supplementary Note). Differentially expressed genes (DEGs, adjusted-p < 0.05) were used for downstream analyses (see GO Semantic Similarity below). In addition to UMAP, we also performed t-distributed Stochastic Neighbor Embedding (t-SNE; Supplementary FIG. S7B) and Principal Component Analysis (PCA; Supplementary FIG. S7C) using the Seurat^143^ R package.

### RNA bulk transcriptome sequencing

RNA-seq libraries were prepared using 200 ng of total RNA and the NEBNext rRNA Depletion Kit (NEB, Cat: E6310X), per manufacturer’s instructions. This kit employs an RNaseH-based method to deplete both cytoplasmic (5S rRNA, 5.8S rRNA, 18S rRNA and 28S rRNA) and mitochondrial ribosomal RNA (12S rRNA and 16S rRNA). The mRNA was enriched via poly-A-selection using oligoDT beads and then the RNA was thermally fragmented and converted to cDNA. The cDNA was adenylated for adaptor ligation and PCR amplified. The resulting libraries were quantified using a Qubit fluorometer (ThermoFisher), Bioanalyzer 2100 (Agilent) for library profile assessment, and qPCR (Kapa Biosciences Cat: KK4622) to validate ligated material, according to the manufacturer’s instructions. The libraries were sequenced using the NovaSeq 6000 with 150 bp paired-end reads. RTA (version 2.4.11; Illumina) was used for base calling and MultiQC (version 1.7) for quality control. Reads were aligned using STAR^145^ (version 2.5.2b) with default parameters to the STAR hg38 reference genome. Gene counts were obtained using the featureCounts^146^ package (version 1.6.4) within the Subread package. The gene transfer format (GTF) file for the genes analyzed in the sc-RNA-seq data (provided by 10X Genomics and used in the *Cell Ranger* pipeline, generated from the hg38 reference transcriptome) was used to better facilitate internal comparison between sc-RNA-seq and bulk RNA-seq datasets.

### RNA bulk transcriptome sequencing analysis

RNA-seq data was analyzed using the DESeq2^147^ R package. Counts were transformed using the regularized logarithm (rlog) normalization algorithm. PCA was performed using the *prcomp* function in R (Supplementary FIG. S8A) and hierarchical clustering using the *hclust* R function with a Ward’s minimum variance method (Supplementary FIG. S8B). Data was visualized using the *ggplot2* R package.

### GO semantic similarity analysis

Genes associated with unique IMPACT mutations (classified as *low*, *moderate*, or *high* IMPACT scores, see above) were identified for each comparison set (i.e., cell line versions, sublines, or sublines including DS8). Additionally, DEGs from the sc-RNA-seq statistical comparisons for each comparison set were determined (see above). The two gene lists were independently subjected to a GO enrichment analysis using *EnrichR*^148^ (version 2.1). Genes were compared to ontology databases *GO Biological Process 2018* and *GO Molecular Function 2018*. The GO terms significantly enriched in the IMPACT mutations (p < 0.05) and in DEGs (adjusted-p < 0.05) were identified and stored independently as separate GO term lists. The significant GO term lists were further separated into GO types, which created GO term lists unique for each combination of sample (cell line version or subline), dataset (IMPACT mutations or DEGs), and GO type (“Biological Process” or “Molecular Function”). For each sample, the mutation and DEG GO term lists associated with each GO type were compared using the semantic similarity metric from Wang et. al.^104^, as calculated in the *GOSemSim* package^149^. This approach compares two individual GO terms using the underlying GO term network structure (see Supplementary Note). Pairwise similarities were calculated on the lists of terms to generate similarity matrices for each sample (e.g., Supplementary Table S2). These similarity matrices were summarized into a single number using the best match average (BMA) method (default in *GOSemSim*; see Supplementary Note). The single metric provides a relative score (between zero and one) that indicates how well the significant GO terms from the IMPACT mutations agree with the significant GO terms from the DEGs. Intuitively, the similarity score quantifies how predictive *differences* in the genomic states of samples (cell line versions, sublines) are of *differences* in the transcriptomic states, with high scores suggesting a genetic basis for the differences and low scores indicating an epigenetic explanation, i.e., molecular differences between basins within a common epigenetic landscape, in accordance with the G/E/S heterogeneity framework (FIG. 1A). Data were visualized using the *ggplot2* R package.

### *In silico* modeling of clonal Fractional Proliferation

#### Birth-death population growth models

Mathematical models of population growth dynamics were constructed using PySB^150^, a Python-based kinetic modeling and simulation framework. We modeled cell proliferation as a simple birth-death process,

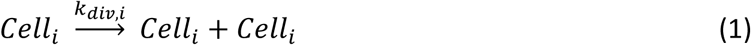

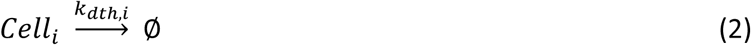

where *i* is an integer index, *k_div,i_* and *k_dth,i_* are division and death rate constants, respectively, for cell type *i*, and ∅ denotes cell death. Note that there is no state switching included in the model. Models with one cell type were used to compare against experimental cFP data for the majority of the PC9-VU sublines (DS1, DS3, DS4, DS6, DS7, DS9), while a two-cell-type model was used in one case (DS8; see Supplementary Note for additional details).

#### Stochastic simulations and in silico DIP rate distributions

All model simulations were run using the stochastic simulation algorithm^73, 107^ (SSA; see Supplementary Note), as implemented in BioNetGen^151^ (invoked from within PySB), to capture the effects of random fluctuations in division and death on cell population proliferation. We performed *in silico* cFP assays, where numerous single cells (run as independent simulations) were grown into colonies of variable size over eight days of simulated time using the SSA and fixed rate constants for division and death (*k_div_* = 0.04×ln(2) h^-1^, *k_dth_* = 0.005×ln(2) h^-1^; Supplementary Table S3), based on vehicle-control proliferation data (Supplementary FIG. S3C; in the case of two cell types, both types were assumed to grow at the same rate outside drug). We ran as many simulations as there were experimental cFP trajectories for the PC9-VU subline being compared against (FIG. 5A and Supplementary FIGS. S9A and S10E). Drug treatment was then modeled by changing the rate constants for division and death (for two cell types, each was assumed to proliferate at different rates in drug; Supplementary Table S3) and running for an additional nine days of simulated time. Simulated time courses were plotted at the same time points as in the corresponding experimental cFP assays for direct comparison (FIG. 5B and Supplementary FIGS. S9B and S10E). *In silico* DIP rates were obtained by taking log_2_ of the total cell counts and calculating the slope of a linear fit to the time course from the time of drug addition to the end of the simulation using the SciPy^152^ *linregress* function. DIP rates for all *in silico* colonies were compiled into a distribution and compared to the corresponding experimental cFP distribution using the Kolmogorov-Smirnov (K-S) test^153^ (FIG. 5C and 6F and Supplementary FIG. S9C).

#### Parameter scans

To account for fluctuations in rates of division and death due to fluctuations in molecular concentrations within and across cells within a population, we repeated the simulation procedure above over ranges of physiologically plausible division and death rate constant values (Supplementary Table S3). For each parameter set (either one or two pairs of post-drug division/death rate constants, depending on subline), we plotted p-values from K-S tests for simulated vs. experimental DIP rate distributions in a heatmap using the *ggplot2* R package (FIGS. 5D and 6G and Supplementary FIG. S9D). We defined a threshold of p<0.1 for statistical significance (i.e., the null hypothesis that the DIP rates are drawn from the same distribution can be rejected) and set the shade of the heatmap element on a graded scale between 0.1 and 1 (the degree of shading has no statistical significance, it is for visualization purposes only). Thus, all colored elements in the heatmap represent sets of division/death rate constants that produce DIP rate distributions statistically indistinguishable from the experimental distributions obtained from cFP assays. Note that the scan for the two-cell-type model (DS8) was performed in four dimensions (two division and two death rate constants) but results were plotted in two dimensions for visual simplicity (FIG. 6G).

## Model and Experimental Analysis Code Availability

The codes used to generate model simulations and analyze experimental data are publicly available via GitHub (github.com/QuLab-VU/GES_2020), or from the corresponding author upon request.

## Data Availability

The sequencing datasets generated in this study can be found in the gene expression omnibus (GEO; GSE150084) and sequence read archive (SRA; PRJNA631050 and PRJNA632351). Additional experimental data will be available from the corresponding author upon request.

## ACKNOWLEDGEMENTS

This work is dedicated to the memory of our friend and colleague Melaine N. Sebastian. We thank Jing Hao for reagent acquisition and Tony Capra, Bishal Paudel, Christian Meyer, Sarah Maddox Groves, Carlos Lopez, Alissa Weaver, John McLean, and Ken Lau for useful discussions.

Sequencing studies were performed by the Vanderbilt Technologies for Advanced Genomics (VANTAGE, Vanderbilt University Medical Center), an institutionally-supported core facility with help from Angela Jones, Karen Beeri, Jamie Roberson, Latha Raju, and Matthew Scholz. Sorting of labeled cells and single-cell seeding for cFP assays were performed with oversight from the Flow Cytometry Shared Resource (Vanderbilt University Medical Center). Drug changes on 384-well plates for cFP assays were performed using the Vanderbilt HTS Core facility. Data processing and model simulations were performed using Advanced Computing Center for Research and Education (ACCRE, Vanderbilt University).

This work was supported by the following funding sources: C.E.H., National Institutes of Health (NIH) Ruth L. Kirschstein National Research Service Award (NRSA, F31-CA221147) and Chemical-Biology Interface Training Grant (T32-GM0650); L.A.H., Vanderbilt Biomedical Informatics Training Program (NLM 5T15-LM007450-14) and Quantitative Systems Biology Center at Vanderbilt; D.R.T., Lung Cancer Research Foundation (LCRF, UALC 13020513) and NIH Research Specialist Award (1R50CA243783); P.L.F., NIH NRSA (F31-CA165840); C.J.R., Vanderbilt Trans-Institutional Programs Grant: Understanding the Complexity of Life One Cell at a Time; V.Q., NIH Clinical and Translational Science Award (U54-CA113007); Sequencing studies were supported by the Vanderbilt Institute for Clinical and Translational Research (VICTR, Voucher VR52385).

## AUTHOR CONTRIBUTIONS

Conceptualization, C.E.H., D.R.T., L.A.H., and V.Q.; Investigation and experiments, C.E.H., C.J.R., D.R.T., and P.L.F.; Bioinformatic Analysis, C.E.H. and C.J.R.; Modeling, C.E.H. and L.A.H.; Writing, C.E.H. and L.A.H.; Review and Editing, C.E.H., D.R.T., L.A.H., and V.Q.

## SUPPLEMENTARY INFORMATION

### SUPPLEMENTARY NOTE

#### Building an *in vitro* model system to mimic tumor heterogeneity

To provide experimental evidence for the G/E/S heterogeneity framework, an experimental system was needed that could explain both an independent level of heterogeneity and other levels dependent upon it (if necessary). For example, if a cell population is genetically distinct, the experimental system must have the capacity to explain genetic differences but also the underlying epigenetic heterogeneity in that cell population and the stochastic variability in those epigenetic basins. For genetically identical but epigenetically distinct cell populations, the framework must be able to explain the epigenetic differences between them and stochastic variability within each epigenetic basin. For cells that are genetically identical and epigenetically identical (i.e., cells derived from an expanded clone), only stochastic variability must be explained. With this idea in mind, we aimed to generate a *family* of cell line ‘versions’ and ‘sublines’ that could capture different levels of heterogeneity. This cell line family can be viewed as a proxy for a heterogeneous tumor composed of multiple genetic states, each of which has multiple epigenetic states, within which stochastic fluctuations occur and occasionally drive phenotypic state changes.

For the cell line family, our goal was to minimize the extent of genetic variability up front, so as not to complicate the potential range of genetic variability that could be identified. Different cancers^1, 2^, driver oncogenes^3–5^, and patient cohorts^6, 7^ exhibit varying degrees of genetic variability, so much so that the field of cancer heterogeneity was originally pioneered just based on genetic variations. However, we limited our study to just genetic variability that arose from a recent common ancestor derived from a single patient with non-small cell lung cancer (NSCLC) and a driver mutation in the epidermal growth factor receptor (EGFR) gene. Therefore, the extent of our genetic heterogeneity exploration was restrained to the recent progression of mutations, specifically those that led to drug-response variability. We also needed a system that could identify non-genetic variability, specifically epigenetic variability in a single common genetic background. Although non-genetic variability is a recent addition to the study of cancer heterogeneity^8^, it has been shown to have distinct and diverse impacts on phenotype^9–11^. Therefore, we aimed to find a system with distinct epigenetic basins that exhibit variable drug-response phenotypes. Finally, we sought to identify non-genetic stochastic variability in clones derived from single epigenetic basins. Previous work has shown that stochastic state transitions can drive daughter cells derived from the same parent to exhibit distinct phenotypes^12–14^. Here, we sought to show that stochasticity alone can lead to some variability in drug-response phenotypes.

Based on the restrictions imposed above and resources available in the laboratory, NSCLC cell line PC9^12, 15, 16^ was identified as a promising candidate. In addition to fulfilling the requirements above, PC9 has a mutation (exon 19 deletion) in the EGFR gene that makes it sensitive to EGFR inhibitors (EGFRi) ^17, 18^, a common treatment for NSCLC. Consequently, response to EGFR inhibition was a natural way to identify phenotypic changes, which we measure in terms of drug response to EGFRi erlotinib. We first identified two versions of the PC9 cell line cultivated at different academic institutions (Supplementary FIG. S2; Massachusetts General Hospital– MGH^12, 16, 18^; Vanderbilt University–VU^19, 20^) in an effort to capture multiple levels of heterogeneity (genetic, epigenetic, and stochastic) in the family. We felt the extended period of separate evolution in cell culture likely allowed for extensive genomic evolution. Therefore, comparison between these two cell line versions allowed for the potential identification of novel genetic, as well as epigenetic, features that differentiate phenotype. PC9-BR1, which had been generated from PC9-VU through dose-escalation therapy in the EGFRi afatinib and became resistant to EGFR inhibition^19^, would serve as a positive control for genetic heterogeneity. We next isolated multiple single-cell derived clones of cell line version PC9-VU, which we call “discrete sublines”^21^ (DS). These sublines were derived to avoid genetic heterogeneity completely, by starting from a single clone and therefore a single genetic background, that of PC9-VU. Additionally, the sublines were meant to identify individual epigenetic basins in the underlying PC9-VU epigenetic landscape. Variability within the sublines themselves were used to understand stochastic variability. Since the sublines were (presumably) genetically and epigenetically identical, the only way to explain variability in our framework was through extrinsic or intrinsic stochastic variability. In this way, we built a system designed to capture multiple levels of heterogeneity: genetic, epigenetic, and stochastic.

#### Definition of phenotype using the DIP rate

We previously described the drug-induced proliferation (DIP) rate as an unbiased metric to quantify drug effect in vitro^20^. This metric, which measures the stable cell proliferation rate of a cell population achieved during prolonged treatment, quantifies cellular fitness in a way that can be tied to phenotype. However, the metric is neutral to the molecular properties that differentiate populations and define fitness. Therefore, we needed to leverage the DIP rate as to better differentiate types of heterogeneity. To supplement the standard DIP rate assay (indicated hereafter as population-level; FIG. 2A,C; Supplementary FIGS. S3A,C, S4A,B), we developed a single-cell DIP rate assay. Originally published as the clonal fractional proliferation (cFP) assay^21^, this approach tracks the DIP rates of many single-cell derived colonies of a cell population to generate a distribution of rates. We modified this method slightly from the original publication by flow-sorting single cells into individual wells in a multi-well plate (384-well), as opposed to having multiple colonies in a single well of a plate. This modification removes some of the subjective aspects of the assay, as well as any influence of other cell types (microenvironment, signaling, etc.) on clonal proliferation. The resulting single-cell distribution of DIP rates, specifically the variance of the distribution, shows the extent of drug-response heterogeneity present in a population (FIGS. 2B,D, 5A,C, 6F and Supplementary FIGS. S3B,D,E, S4A,C, S9A,C, S10E). Conversely, the population-level assay provides a single DIP rate, which is reflective of the proliferation rate of the most fit clones in the distribution.

Considering the types of information that can be conferred by the different assays, different types of heterogeneity (and variability) can be broadly distinguished using a combination of the methods. In the case of a single molecular state (same genetic and epigenetic profiles), the population would have a single DIP rate and a narrow DIP rate distribution, reflective of only stochastic noise in the drug-response dynamics. Conversely, a heterogeneous cell population, composed of multiple genetic or epigenetic states, would exhibit a broader (possibly multi-modal) DIP rate distribution, but still a single DIP rate reflective of the most fit states. However, distinguishing between molecular states resulting from genetic versus epigenetic heterogeneity is more difficult. In general, it is not difficult to imagine that genetically identical, epigenetically distinct cell populations (i.e., single-cell derived clones, basins in the same epigenetic landscape) should each have narrow unimodal DIP rate distributions that fall within the broad parental (i.e., entire landscape) distribution. In this case, it would be concluded that the parent cell line exhibits only epigenetic heterogeneity, as seen by the clones. But, in the case of genetically and epigenetically distinct cell populations, the population may still exhibit a broad DIP rate distribution and single DIP rate reflective of the most fit cells. However, epigenetically distinct clones of each genetic background are less likely to overlap with clones of other genetic backgrounds, due to the inherent fitness advantage of *genetic* instead of *genomic* mutations (see next section). Therefore, clones that differ epigenetically are more likely to exhibit phenotypic variability such that their DIP rate distributions are closer to one another, while clones that differ genetically are more likely to have distinct DIP rate distributions. However, the subtlety of these differences necessitates the need for other types of data (genomic, transcriptomic, etc.) to better characterize cell populations, one of the bases for this paper. We address these topics below.

#### Distinguishing genomics from genetics

The following issues related to mutational events and their predicted impact on phenotype should be considered to distinguish events that lead to genetic heterogeneity. In short, not all mutational events generate genetic heterogeneity, insofar as a phenotypic impact.

##### Phenotypic context of mutational events

Mutational events do not all have the same phenotypic consequence. Depending on the phenotype measured, mutations have a contextual dependence. In these cases, we seek to clarify the terminology (see Supplementary Table S1).

Consider a case where two genomes exhibit extensive mutations that distinguish one from the other. In this example, the measured phenotype is the response to drug treatment. However, these mutations fall in regions that do not have an effect on the drug response (functional genomic regions, see Supplementary FIG. S1A). Although this case seems specific, it is actually rather ubiquitous. Among the tens of thousands of coding genes in the human genome, multiple orders of magnitude fewer have been heavily implicated in cancer drug resistance^5^. This means that mutations in a large number of genes will have a negligible effect on phenotype. In this case, based on our terminology (Supplementary Table S1), these two genomes would be considered *genetically identical,* but *genomically distinct*. We use this language because the term ‘genomics’ includes the universe of all potential DNA variations, but ‘genetics’ has implications on heredity and function. Since the mutations that differentiate populations in this case do not fall in functional genomic regions, we do not consider these genomic variations to encode for genetic differences.

Conversely, consider the case where two genomes have widespread mutational differences that do fall in functional genomic regions. In this case, these genomes would be both *genomically distinct* and *genetically distinct*. Depending on the phenotype observed, there may be many or few of these key regions. In the case of cancer, mutations in these functional genomic regions aid survival in a specific treatment^5^. Additionally, functional genomic regions have been shown to be different for distinct drugs and cancer types. The work in this manuscript provides a practical example in which the differentiation between mutations depending on phenotypic context does play a role on phenotype. Therefore, we use the terms *genetic* and *genomic* to distinguish mutational context, and to be consistent with the G/E/S theoretical framework.

##### Differential nature of mutational events

Mutational events, by definition, are changes in the structure of a gene. However, these changes can take many different forms. The first scheme to categorize mutations is *variant classification*. Variant classification falls into two primary forms, sequence variants and structural variants. Sequence variants pertain to mutations of one or a few nucleotides, while structural variants deal with modification of larger genomic regions. In this study, we focus on the sequence variants (examples in Supplementary FIG. S1B-D). Within the sequence variant category, there is a further breakdown into substitutions and indels. Substitutions are alterations where the length of the variant change is the same as the reference region (Supplementary FIG. S1B). Substitutions can occur in a range of nucleotide lengths, but most occur as single nucleotide substitutions, called single nucleotide polymorphisms (SNPs). SNPs are the most common form of sequence variation, and heavily outweigh other length substitutions. The other type of sequence variation, indels, are broken down into insertions, deletions, and combinations (also referred to as indels) (Supplementary FIG. S1C). As the name suggests, insertions are an insertion of one or many nucleotides, while deletions are a deletion of one or several nucleotides.

A second categorization scheme is *variant consequence*. Variants of any classification can encode for any consequence (Supplementary FIG. S1D). Variant consequence can initially be separated into synonymous and nonsynonymous mutations. Synonymous mutations, otherwise known as silent mutations, are changes in the DNA structure that encode for the same amino acid as the reference versions of the gene. Therefore, these genes are considered to have *no impact on phenotype*. Nonsynonymous mutations, conversely, are the class of all other mutations that encode for a different amino acid than the reference gene. These mutations *can have an impact on phenotype*, depending on the type of mutation and aforementioned phenotypic context. Nonsynonymous mutations can be further broken down into missense and nonsense mutations. Missense mutations encode for a different amino acid compared to the reference, while nonsense mutations encode specifically for a start or stop codon where one was not encoded in the reference. Missense mutations can be further subdivided into conservative (same amino acid type – e.g. polar) and non-conservative (different amino acid type – e.g. polar changing to non-polar). In terms of impact, the general rankings of variant consequence (from low to high) is: synonymous/silent, conservative missense, non-conservative missense, and nonsense mutations (Supplementary FIG. S1D). But, depending on the phenotypic context, these rankings *can* be variable. For example, the canonical EGFR T790M mutation that encodes for resistance to EGFRi, is a missense mutation that has profound impact (FIG. 3E).

##### Determination and quantification of IMPACT mutations

Defining the consequence of all possible mutations in cancer is intractable, and likely not particularly useful^5^. This is primarily because mutations can have an impact on various regions of molecular networks not directly adjacent to the mutated gene. In this paper, we have made an attempt to extend the scope of key mutations driving genetic heterogeneity in the PC9 cell line family. Here, we have used classifiers defined by the *Ensembl* Variant Effect Predictor (VEP)^22^ to better understand phenotypic effect. To this end, we utilized the VEP *IMPACT* score to rate the predicted effect of individual mutations. The IMPACT score is a subjective classification of mutational severity based on mutation classification, consequence, and predicted protein structure impact, similar to other tools like SnpEff^23^. Therefore, the IMPACT classifier helps narrow the entire field of mutations to ones more likely to have a phenotypic effect. We used the IMPACT score to determine the total number and relative proportion (compared to the number of unique mutations in each sample; FIGS. 3C, 4C, 6B) of consequential mutations, with the rationale that more IMPACT mutations lead to a greater chance that a single mutation may initiate *genetic heterogeneity*. This score was utilized to establish a baseline number of consequential mutations (FIG. 3C) by which to compare sublines (FIGS. 4C, 6D), and played a significant role in the determination that DS8 is genetically distinct.

##### Using nonsynonymous mutational load to predict distinguishing mutations

An additional layer of mutational significance was utilized to identify gene sets that differentiate cell populations. We used *dNdScv*^24^, a method that compares the nonsynonymous and synonymous mutational of individual genes to identify genes under positive selection (see Methods). Similar to other methods that quantify mutational significance^5^, dNdScv uses a statistical model to predict genes that have a high relative nonsynonymous mutational load and therefore could be phenotypically significant (e.g., mutations enriched in a cell population resistant to treatment). In our case, we used this method to identify a list of candidate genes in which the mutational load could distinguish samples within our comparison groups (FIG. 3B - cell line versions, FIG. 4B – sublines without DS8, FIG. 6A – all sublines). This metric confirmed the large number phenotypically significant genes mutated in cell line versions (specifically PC9-BR1, FIG. 3D) and the fewer mutations by the same signature in sublines (FIG. 4D, Supplementary FIG. S10B). In principle, methods like these can be utilized to determine mutations that drive phenotype.

#### Connecting transcriptomics to epigenetics

Like mentioned in the main text, the definition of epigenetics used in the context of this paper is derived from the traditional definition, which roughly translates to anything other than genetics^25^. Ergo, the epigenetic landscape we present here is a theoretical construct that attempts to encapsulate that definition of epigenetics. Some researchers have attempted to put a mathematical formalism to this definition of epigenetics^26, 27^, with variable success. Contrary to expectation, the best way to currently measure the epigenetic landscape is with gene expression data, specifically sc-RNA-seq. The rationale for this conclusion has several components. First, since gene expression is a count of all measurable genes, it is considered to be a snapshot of the molecular network at a point in time. Because the steady-states of the network encode for basins in the epigenetic landscape, gene expression information is necessary to construct the landscape, although not limited to just that information, in principle. Second, a single-cell approach provides some insight into transition rate probabilities between epigenetic states, another feature of the landscape^28^. The number of cells that have a similar transcriptomic signature can be used to calculate landscape features such as depth of the well and barrier heights, as well as predict potential transition events. Other omics-based data, such as epigenomics, proteomics, and metabolomics, could be considered as additional reflections of the landscape (not genetics because it establishes the landscape, see next section for details). Unfortunately, none of these approaches have been developed at the single-cell level with high quality, genome-wide coverage like sc-RNA-seq. Also, although it may seem like epigenomics may be the best way to calculate the epigenetic landscape, approaches like ATAC-seq and ChIP-seq (i.e., epigenomics assays) only capture one or a few types of epigenetic marks (e.g. chromatin accessibility, methylation, etc.), and therefore do not capture the breadth of information necessary to calculate an entire landscape. In principle, an epigenetic landscape could be calculated from most types of measurable epigenetic information. However, it would lack the information necessary to understand the entire molecular state of a cell, one of the defining factors of the epigenetic landscape.

#### The genetic-to-epigenetic connection

In the context of the G/E/S heterogeneity framework, genetics are fundamentally tied to epigenetics. The genetic state encodes for a complex biochemical network that determines the epigenetic landscape. Significant research has been put into this idea, only to find that the connection does not exhibit a 1:1 correlation^27^. Due to this problem, it is virtually impossible to perform a gene-by-gene comparison of genomics (encodes for genetics) to transcriptomics (encodes for epigenetics) data that provides rational information. Instead, we aimed to develop a broader way to identify this connection. In the main text of the paper, we make several claims with different types of data that support the idea that populations are genetically and/or epigenetically distinct. However, these claims were made at the level of one measurable phenotype (drug response, genomics, transcriptomics). We wanted to supplement this data with a metric that connects genetics to epigenetics, but not at the level of single genes (for reasons mentioned above). We settled on the *process* level, which we attempted to reflect in gene ontology (GO) term similarity.

GO terms are structured as directed acyclic graphs, composed of nodes and edges^29, 30^. This graph-based structure is made to model pairwise relationships between objects, in this case GO terms, where the terms are the nodes and relationships between terms are the edges. These GO terms broadly fall into levels, with the lowest number level (one) reflective of the most general term and the highest-level term (variable) as the most specific. For example, in the case of the cellular component ontology type, the level one term would be “cellular component,” and an edge connects to “cell,” followed by “intracellular,” “intracellular organelle,” and so on. The complexity of these graphs can be immense and enriched GO terms often arise from different levels, making direct comparisons between individual GO terms largely ineffective.

For these reasons, we decided to use a score that compares GO terms as node-edge graphs. Specifically, we utilize *GoSemSim*^31^, an R-based tool to calculate GO term semantic similarity. We employed the implementation developed by Wang et al.^32^ that compares the GO term graph structure. In this implementation, a similarity score is returned for the comparison of two GO terms. This score is calculated by comparing the similarity between subsets of the ontology graph associated with the individual terms and their upstream (lower level) terms. In our case, we generated two lists of GO terms for each sample, one from the genomics (unique genes with IMPACT mutations) and one from transcriptomics (differentially expressed genes from sc-RNA-seq) data (see Methods). We fed these genes into independent GO enrichment analyses to get two lists of these GO terms that were statistically enriched for each data type (genomics and transcriptomics). We then calculated the semantic similarity between the genomics- and transcriptomics-defined statistically significant GO term lists, which generated a matrix of similarity scores (example matrix in Supplementary Table S2). We used a modified averaging metric (Best Match Average (BMA) - recommended by the authors of GoSemSim^31^) to generate a single similarity term between the genomics and transcriptomics for each cell population (FIGS. 3I, 4I, 6E). The BMA metric identifies the maximum semantic similarity score for each row and column in the aforementioned similarity matrix, and averages those maxima to determine the final score. Using this approach, scores are not heavily weighted by the other similarities in the matrix, many of which are low and skew the differences between GO term lists. In this way, we achieved an indication for the amount of influence genetics had on epigenetics in a single cell line version or subline, with the expectation that populations with a larger genetic influence would exhibit a higher score than those that did not.

#### Simple growth model of stochastic birth and death

##### Monoclonal growth model and connection to DIP rate

Biological noise, often referred to as stochasticity, can be broken down into extrinsic and intrinsic components. Extrinsic noise pertains to variations in the environment, while intrinsic noise refers probabilistic division and death events that results in fluctuations, especially at low cell numbers^33, 34^. Many models of cell population dynamics have simulated drug response dynamics with extrinsic noise, using deterministic simulations with various factors influencing cellular dynamics (initial conditions, mutation rates, etc.)^35, 36^. However, few (especially in cancer) have studied the intrinsic noise associated with cell fate decisions (i.e., division and death). We previously described the drug-induced proliferation (DIP) rate, which was derived using a simple mathematical model of cell division and death^20^. Here, we extend that framework to include intrinsic stochastic variation using the Gillespie stochastic simulation algorithm (SSA)^37–40^. The SSA was specifically employed to test whether the DIP rate variability seen in subline clonal fractional proliferation (cFP) assays were indeed due to fluctuations in single epigenetic basins (i.e., sublines).

To generate a model of stochastic cell population dynamics, we started with a simple model of cell division and death, encoded by the following chemical reactions:

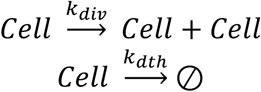

These reactions encode for the ordinary differential equation (ODE):

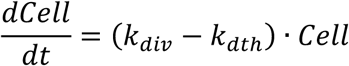

In a deterministic solution, the equation above would be solved over individual time steps to simulate a trajectory. However, since we are simulating this model stochastically, we use a different method to generate a solution. Specifically, we aim to solve the chemical master equation (CME), the general form of which is:

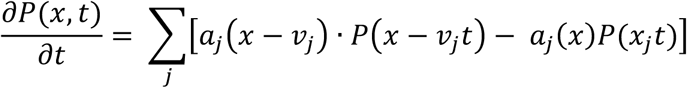

where **P(x,t)** is the probability of being is state **x** in time **t**, **a_j_** is the probability that a reaction will occur sometime between time **t** and the next infinitesimal stochastic time step, and **v** is the state change vector. Generally, the equation determines the probability that a molecular state will have a certain population at a future point in time by calculating the flux into and out of that state. In the case of our MGM, the equation would be calculated:

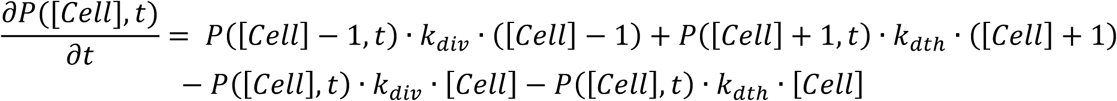

where **[Cell]** is the number of cells at any point in time **t**. As follows, this system generates a system of coupled ODEs, which generally cannot be solved analytically. Many attempts have been made to approximate the CME, such as reducing the state space with the finite state projection method^41^. Here, we use the Gillespie SSA^37–40^, which works off the fundamental assumption that a reaction of a certain type will fire randomly, but proportionally to the reaction rate. Specifically, stochasticity is generated from two randomly drawn numbers. The first random number (r_1_) is drawn from a uniform distribution in the unit interval to determine **τ**, the stochastic time step:

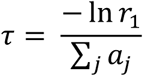

The second random number (r_2_) is drawn proportionally to the reaction rate, where:

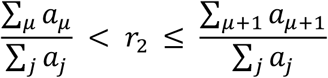

so that r_2_ will direct the μ+1^th^ reaction to fire (j is the final reaction). However, the proportionality of these reactions is determined by the reaction rate, which is a function of the reaction rate constant (*k_div_* and *k_death_* here). This proportionality is broken down in the term **a_μ_**, which is calculated:

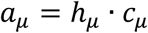

where **h_μ_** is the number of reaction combinations and **c_μ_** is the reaction rate constant for the **μ^th^** reaction. In our case, we are simulating a first-order reaction, so **h_μ_**=1. **c_μ_** is the summation of all the reaction rate constants, making it *k_div_ + k_dth_*. **a_j_** is a function of the concentration of state **x** at any given time **t**, so in the case of the model:

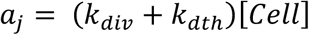

So, r_1_ will determine **τ** and r_2_ will determine the decision for a simulated cell to divide or die (based on **a**). The system then updates, and the algorithm determines whether to stop or continue proceeding (called a check-stop). If determined to continue, it will redefine the reaction rate, recalculate the random numbers, and simulate a result again.

Together, we call this the monoclonal growth model (MGM) because it is meant to reflect the growth dynamics of a single clone (or clonal-derived subline). More specifically, the MGM was designed to simulate the stochastic growth of cells residing in a single basin in the epigenetic landscape, making these cells both genetically and epigenetically identical, hence the single division and death rates. Therefore, the only source of variability expected in these cells would be stochastic fluctuations. As noted in the main text and above, this approach was used to explain the DIP rate distribution of single-cell derived clonal sublines (FIG. 5), with the exception of DS8 (see below). In an effort to exhaustively simulate the parameter space, we ran a large parameter scan across a wide array of division and death rates, with minimal spacing between parameter combinations (FIG. 5D and Supplementary FIG. S9D). The scan identifies many division and death rate combinations where the simulated distribution cannot be statistically distinguished from experimental subline distributions, showing that the model is not overly sensitive to chosen parameters. Importantly, it also indicates that fluctuations in molecular state (i.e., small differences in the division and death rate constants) can vary within epigenetic basins.

##### The polyclonal growth model

In the case of DS8, the MGM (one division and death rate plus stochasticity) could not explain the variability in the DIP rate distribution, specifically the bimodal distribution. So, we generalized the one-state model into multiple states, which we call the polyclonal growth model (PGM). The PGM is an extension of the MGM to include multiple cell types, each with different division and death rate combinations. Therefore, the PGM is identical to the MGM shown above, but just multiple MGMs in parallel.

To explain the DS8 DIP rate distribution, we first attempted a two-state model, which consisted of two sets of division and death rates (four total rate parameters). In order to generate the DS8 bimodal distribution, the underlying DIP rates (i.e., division minus death rate) needed to be appreciably different, so we sampled distinct regions of parameter space for each state, corresponding to the DIP rates underlying each mode of the DS8 distribution. We performed a four-dimensional parameter scan (two division, two death rates), and projected into two dimensions, where two discrete states could be identified (FIG. 6F) based on concordance between the experimental DS8 and simulated PGM distributions. Similar to the MGM, the PGM parameter scan identifies many division and death rates that result in DIP rate distributions that statistically agree with experimental results (FIG. 6G). Although the PGM is agnostic to how the population achieved the two states (i.e., genetic or epigenetic, see below), this is another example of fluctuations in molecular state due to intrinsic stochasticity.

A natural extension of the PGM would be to include transitions between cell states, but we do not include them here to simplify the model and results. Future studies will aim to incorporate models that simulate PGMs with transitions (i.e., epigenetic landscapes), each with multiple states (i.e., epigenetic basins).

#### Justification for multiple genetic states in DS8

As noted in FIG. 7, DS8 seems to have evolved a genetic mutation that made it nearly resistant to EGFRi. This process could have happened two different ways. First, DS8 could have acquired a key genetic mutation after single-cell cloning, meaning that both the PC9-VU and a new genetic state exist in DS8 (FIG. 7, Supplementary FIG. S10A). In this case, each genetic state would have an epigenetic landscape and therefore multiple sets of epigenetic basins available. Another possibility is that DS8 could have genetically evolved prior to single-cell isolation as a single genetically distinct clone within PC9-VU (Supplementary FIG. S10B). In this case, the DS8 bimodal DIP rate distribution is reflective of two (or more) epigenetic basins in a single epigenetic landscape (i.e., one common genetic background). We believe that the former option is the case. The similarity of the DS8 left distribution with all other PC9-VU sublines suggests that the PC9-VU epigenetic landscape is present in DS8. Alternatively, in the latter case, the new DS8 epigenetic landscape would likely be different from the PC9-VU epigenetic landscape (because genetic mutations change the underlying biochemical network), leading to distinct division and death rates and DIP rate profiles. Although it is possible that DS8 could have acquired a mutation that resulted in an epigenetic landscape similar to PC9-VU, it is unlikely that a similar network would have such a distinctly different DIP rate in EGFRi (FIG. 2C). Additionally, a new state is clearly present in DS8 as the right mode of the DIP rate distribution, which has no overlap with any subline DIP rate distribution, and minimal with the PC9-VU DIP rate distributions, lying predominately between PC9-VU and PC9-BR1. This is consistent with a new genetic state that was not present in PC9-VU or sublines.

Therefore, we believe that DS8 encapsulates two genetic states, one from PC9-VU and another new, each with epigenetic landscapes. In these landscapes, a large proportion of cells likely lie in a single epigenetic basin, but potentially multiple basins theoretically exist and would be available for future epigenetic state transitions (driven by stochastic fluctuations). As DS8 expanded (drug-naïve), both the sensitive and non-sensitive genetic states co-existed, with likely similar proliferation rates (since the drug response phenotype is revealed in EGFRi, we cannot know the drug-naïve proliferation rate of each state a priori). At the time of treatment in the population-level DIP rate assay, the more fit genetic state (and its epigenetic basin(s)) was reflected in the DIP rate (FIG. 2C) due to clonal selection. However, in the cFP assay, multiple genetic states were identified because single cells were randomly measured from the population and therefore no clonal selection could skew the DIP rates (FIG. 2D), resulting in the DS8 bimodal DIP rate distribution.

In addition to the PGM, multiple genetic states in DS8 are reflected in its genomic signature. Although DS8 does not show significant genetic heterogeneity as shown by the cell line version genetic signature (Supplementary FIG. S10B), it does have two mutations in cancer-associated genes (Supplementary FIG. S10C), one of which is a missense mutation in ABC5, a component of transmembrane ATP-binding cassette (ABC) transporters, which utilize ATP binding and hydrolysis to transport organic material across membranes^42^. Considering the drug-resistance implications of ABC transporters^43^, it is possible that ABC5 is a key genetic mutation that establishes a new genetic state within DS8. DS8 also exhibits a mutation in the RELN gene, but considering it is synonymous, the G/E/S heterogeneity framework considers it genomic (instead of genetic, see Supplementary Table S1) and therefore it is not predicted to have an effect on phenotype. Interestingly, DS3 also exhibits one unknown mutation type (i.e., cannot be defined by substitution or indel) in DACH1, which has been implicated in tumor growth suppression^44^. However, DS3 showed no clear functional change in terms of the DIP rate (FIG. 2C,D), overall genomic (FIG. 3A-F), or transcriptomic profiles (FIG. 3G,H), as well as no clear genetic-to-epigenetic connection (FIG. 3I), suggesting that the genomic change is not encoding a genetic change. Additionally, a clear distinction in the DS8 genetic state can be seen in our identity-by-state analysis of the SNV profiles (Supplementary FIG. S12A), which shows DS8 in a completely separate region of the reduce dimensionality PCA space. Taken together, the genomic information alone points to clear genetic differences in DS8.

To supplement the model and genetic information that distinguishes DS8, we also identify key transcriptomic differences. First, we see a clear distinction between DS8 and all other cell line versions and sublines using bulk RNA-seq (Supplementary FIG. S8). We then use sc-RNA-seq to determine the proportionality of DS8 cells in a common transcriptomic space (bulk RNA-seq does not capture the proportionality, hence the need for single-cell resolution). For example, do all DS8 cells reside together, like PC9-MGH? Or are they split across different regions of the space? As mentioned in the main text, DS8 mainly occupies a completely distinct region of the single-cell transcriptomic UMAP space (FIG. 6C and Supplementary FIG. S7A), as well as in other dimensionality reduction approaches (Supplementary FIG. 7B,C). However, some DS8 cells do lie in regions of the UMAP space occupied by PC9-VU and other sublines, suggesting that DS8 is composed of a genetic state inside PC9-VU (see above). This result provides further confirmation for the proposed hypothesis above, that DS8 is composed of at least two genetic states, one from PC9-VU and another newly acquired after single-cell isolation. Interestingly, DS9 is the only other subline that has cells overlapping with the larger cohort of DS8 cells in the UMAP space, especially considering it exhibited the largest initial death of any subline following EGFRi (FIG. 2C). This small minority of cells are less than 0.1% of the sampled DS9 population, and therefore did not show up appreciably in other analyses (presumably due to lack of single-cell approaches and detection power). However, it is possible that sublines, such as DS9, have small minorities of cells that exhibit genetic differences but do not play a large role in the drug response phenotypes, at least in the short term. Additionally, in line with the postulated views of Huang^45, 46^, it is possible that cell populations that fall into minority regions of the epigenetic landscape may actually lie in close proximity to unoccupied regions of the landscape. These unoccupied regions may be opened by a major event, such as a key genetic mutation, that lowers the activation energy and allows for more probabilistic transitions into the unoccupied regions (Supplementary FIG. S11B). Further information and analysis would need to be performed to determine if that hypothesis is indeed a possibility.

All said, the majority of the evidence points to DS8 being composed of two genetic states, one from PC9-VU and another novel drug-resistant state. However, the discrepancy between a one- vs. two- genetic state DS8 does not modify the findings within the paper or change the underlying heterogeneity framework (Supplementary FIG. S11). Cancer cells are constantly acquiring new genomic mutations, which may or may not lead to functional differences. In the case of DS8, genetic changes were identified and the DIP rate profiles are distinct, leading to the rational conclusion that the genetic changes caused the resistance to EGFRi.

#### The role of stochasticity on the epigenetic landscape

The results of the MGM and PGM show that intrinsic stochasticity alone can explain variability within subline DIP rate distributions. Although variability could always result from a number of hidden factors, our finding that noise alone *can* explain the variability should help change the narrative that every aspect of biological variability in cancer must be genetic. Intrinsic stochasticity both exists, and can have an effect on phenotype.

Evidence in this paper of intrinsic stochastic fluctuations has provided a mechanism by which cells can move between epigenetic basins. Although not explicitly proven here, the work of others suggests cells residing in epigenetic basins in a common landscape exhibit plasticity^47, 48^, meaning that cells can transition between basins reversibly, which we suggest here are due to stochastic fluctuations. Specifically, basins in which PC9-VU sublines reside fall within the wider PC9-VU epigenetic landscape (with the likely exception of DS8) and have the potential to transition between basins. Here, we have captured clones early in the diversification process and therefore cells may not have had enough time to transition between basins, although larger proportional overlap between sublines that cell line versions (FIGS. 3G and 4G) may provide some evidence for this phenomenon. Since the time necessary to diversify between epigenetic basins cannot be known a priori, it is difficult to effectively determine the stability of these basins, which would be reflected in transition events. However, considering the speed of drug-induced state transitions identified in our previous work in BRAF-mutant melanoma^48^, we can extrapolate that the basins in PC9 are relatively deeper and more stable. It is for this reason that we could capture stable drug-response differences between sublines in the first place (FIG. 2C-D), as opposed to all sublines eventually stabilizing into a common DIP rate. It is also the reason we have not captured explicit transition events. There exists a tradeoff between determination of epigenetic basins and identification of epigenetic state transitions: ability to see many of one type inherently means one can see few of the other. In the case of this paper, we could clearly find epigenetic basins, at the expense of observation of explicit state transitions. Provided unlimited time or more precise tools for lineage tracing^49, 50^, we could identify rare, discrete transition events between epigenetic basins. But, provided the theory (FIG. 1) and observations in this paper, we can extrapolate that because of the stochasticity identified within epigenetic basins, there must exist a mechanism where that noise can push cells over the boundary into new basins. Future publications will explore these transitions in systems where plasticity is more prevalent and happens over a shorter time period.

The G/E/S heterogeneity framework provides a clearer understanding of cellular plasticity and survival by detailing a mechanism by which cells can survive treatment. Cells stochastically transition between basins in the epigenetic landscape, and acquire genetic mutations to create new landscapes with different inherent plasticity. Therefore, the modified model of cellular plasticity in response to treatment is as follows: (1) Cells exist and reversibly transition between epigenetic states in the epigenetic landscape in drug-naïve conditions (proportional to the transition rates). These epigenetic basins can have differential fitness in a single drug treatment, and are different depending on the type of drug; (2) Drug treatment changes the landscape so that cells can preferentially transition into different states, some of which have better fitness in the presence of drug, depending on their previous location in the landscape. Cells that survive the treatment serve as a reservoir for potential new genetic mutations, which happen over many cell divisions while cells reside in the drug-tolerant basins; (3) Key genetic mutations open new genetic state(s) that have distinct epigenetic landscapes. The new epigenetic landscapes have epigenetic basins that exhibit greater fitness in the presence of that drug, making the cells insensitive to that treatment.

### SUPPLEMENTARY TABLES

**Supplementary Table S1.** Glossary of terms.

|  |  |
| --- | --- |
| *Heterogeneity* | Differences (genetic, epigenetic, or stochastic) that exist *within* a cell population. |
| *Variability* | Differences (genetic, epigenetic, or stochastic) that exist *across* cell populations. |
| *Phenotype* | An observable property of a cell population, in the absence or in response to environmental perturbations. In the context of this work, our phenotype of interest is response to EGFR inhibition (EGFRi), which we quantify in terms of single-cell transcriptomic features (no drug) and drug-induced proliferation (DIP) rates. Since cell population-level quantities like DIP rate are driven by underlying molecular regulatory networks, we consider these to be two different presentations of the same phenotype. |
| *Genomic state* | The full sequence of nucleotides that comprise a genome. With four types of nucleotides (A, C, T, G) and ~3.2 billion nucleotides per genome, the number of possible nucleotide configurations is massive (effectively infinite, even when excluding non-viable configurations). Therefore, it is exceedingly rare for two cells to reside in the same genomic state. |
| *Functional genomic regions* | Locations in the DNA sequence that encode for genes that impact the phenotype of interest in some quantifiable way. |
| *Genomic variability* | Differences in the genomic-state compositions across two or more cell populations. Since every cell in a population is likely to exist in a unique genomic state, cell populations will always exhibit genomic variability with respect to each other. In this work, we quantify genomic variability in terms of differences in distributions of called variants between populations in bulk whole-exome sequencing data. |
| *Genetic state* | The sequence of nucleotides that comprise only functional genomic regions of the DNA. Because the scope is much more limited, the number of possible genetic states available to a cell is significantly smaller than the number of genomic states, i.e., many cells, up to and including an entire cell population, may share the same genetic state. |
| *Genetic heterogeneity* | The degree to which cells within a population are distributed across different genetic states. |
| *Genetic clone* | A group of cells that all occupy the same genetic state. |
| *Genetic variability* | Differences in the genetic-state compositions of two or more cell populations (the populations may be genetically heterogeneous). Importantly, cell populations that are genetically identical will still display some degree of genomic variability. |
| *Molecular state* | Populations (or concentrations) of all intracellular biochemical species (e.g., mRNAs, proteins, small molecules) associated with a phenotype of interest. Because copy numbers can be large and fluctuate due to intrinsic and extrinsic sources of intracellular noise, it is exceedingly rare for two cells to occupy the same molecular state. |
| *Epigenetic landscape* | A conceptual and mathematical construct that assigns a “quasi-potential energy” to all possible molecular states of a cell. Fundamentally, the epigenetic landscape is the consequence of the complex biochemical interaction networks that underlie the phenotype of interest (i.e., different phenotypes will have different associated landscapes). Altering the molecular species, interactions, or rate parameters (e.g., binding affinities, synthesis and degradation rates) within these networks will change the nature of the landscape. Since many rate parameters depend on the molecular structures of gene products, the epigenetic landscape can be seen as associated with a specific genetic state. Relatedly, mutations that change the structures of gene products change the topography of the associated epigenetic landscape. |
| *Epigenetic state* | Often used interchangeably with “phenotypic state,” a molecular steady state of a cell *and all neighboring molecular states* that tend to gravitate towards it in response to small perturbations. Mathematically, the steady state is known as an “attractor” and the collection of states associated with it a “basin of attraction.” Conceptually, an epigenetic landscape can be seen as a collection of epigenetic states (local minima) separated by barriers, with traversal of these barriers corresponding to phenotypic state transitions. Importantly, cells can differ in their molecular states (e.g., due to fluctuations in gene expression) but still occupy the same epigenetic state. Experimentally, epigenetic states can be revealed by examining *subsets* of the molecular species that define the epigenetic landscape (mathematically, low-dimensional *projections* of the phenotypic state space), e.g., with single-cell transcriptomics. |
| *Epigenetic heterogeneity* | The degree to which cells within a population of *genetically identical* cells are distributed across different epigenetic states within a common epigenetic landscape. |
| *Epigenetic clone* | A group of genetically identical cells that all occupy the same epigenetic state. |
| *Epigenetic variability* | Differences in the epigenetic-state compositions of two or more cell populations (not necessarily genetically identical). Because stochastic fluctuations drive phenotypic state transitions, there is almost always some degree of epigenetic variability between cell populations (an important exception being low-passage number single cell-derived colonies from the same epigenetic state). |
| *Stochastic heterogeneity* | The degree to which cells within a population of genetically and epigenetically identical cells vary across molecular states. |
| *Stochastic variability* | Differences in the molecular-state compositions across two or more genetically and epigenetically identical cell populations arising from fluctuations in gene expression, randomness in cell division, and other sources of biological noise. Importantly, stochastic variability can present as phenotypic differences across epigenetically identical populations, e.g., differences in DIP rates across single cell-derived colonies in clonal Fractional Proliferation assays. |
| *Genetic-to-epigenetic connection* | The degree to which mutations unique to a cell population, with respect to one or more other cell populations, are predictive of epigenetic differences seen among the cell populations. For example, cells occupying two epigenetic states within a common epigenetic landscape are, by definition, genetically identical. Therefore, any *genomic* differences seen between them should have no bearing on epigenetic differences. In practice, however, there is likely to be some correlation, simply by chance, between mutated genes and epigenetic (e.g., transcriptomic) differences. Thus, in this work, we assess the degree of the genetic-to-epigenetic connection of a cell population by comparing against the genetic-to-epigenetic connections of cell populations within a group of populations known to be genetically distinct (the cell line versions). |

**Supplementary Table S2.** Example GO (molecular function) semantic similarity matrix for DS8.

|  | GO:0003723 | GO:0045296 | GO:0004866 | GO:0061135 | GO:0030414 |
| --- | --- | --- | --- | --- | --- |
| GO:0016878 | 0.111 | 0.122 | 0.087 | 0.122 | 0.109 |
| GO:0004321 | 0.084 | 0.092 | 0.065 | 0.092 | 0.083 |
| GO:0003996 | 0.087 | 0.094 | 0.068 | 0.094 | 0.085 |
| GO:0016405 | 0.111 | 0.122 | 0.087 | 0.122 | 0.109 |
| GO:0016408 | 0.096 | 0.105 | 0.074 | 0.105 | 0.094 |
| GO:0042162 | 0.589 | 0.196 | 0.062 | 0.087 | 0.079 |
| GO:0015645 | 0.111 | 0.122 | 0.087 | 0.122 | 0.109 |
| GO:0016820 | 0.103 | 0.111 | 0.084 | 0.111 | 0.102 |
| GO:0015405 | 0.096 | 0.105 | 0.074 | 0.105 | 0.094 |
| GO:0042626 | 0.059 | 0.062 | 0.049 | 0.062 | 0.059 |
| GO:0046966 | 0.198 | 0.36 | 0.071 | 0.094 | 0.086 |
| GO:0043492 | 0.064 | 0.069 | 0.048 | 0.069 | 0.063 |
| GO:0003697 | 0.659 | 0.219 | 0.07 | 0.097 | 0.088 |
| GO:0008191 | 0.067 | 0.072 | 0.798 | 0.56 | 0.65 |
| GO:0015075 | 0.102 | 0.111 | 0.081 | 0.111 | 0.1 |
| GO:0042809 | 0.198 | 0.36 | 0.071 | 0.094 | 0.086 |
| GO:0030676 | 0.096 | 0.105 | 0.167 | 0.235 | 0.211 |
| GO:0010485 | 0.058 | 0.063 | 0.043 | 0.063 | 0.057 |
| GO:0005261 | 0.063 | 0.067 | 0.05 | 0.067 | 0.062 |
| GO:0005158 | 0.209 | 0.383 | 0.074 | 0.1 | 0.091 |
| GO:0030507 | 0.25 | 0.465 | 0.087 | 0.122 | 0.109 |
| GO:0043015 | 0.216 | 0.399 | 0.074 | 0.105 | 0.094 |
| GO:0098641 | 0.182 | 0.713 | 0.064 | 0.087 | 0.079 |
| GO:0016864 | 0.111 | 0.122 | 0.087 | 0.122 | 0.109 |
| GO:0003756 | 0.096 | 0.105 | 0.074 | 0.105 | 0.094 |
| GO:0035254 | 0.25 | 0.465 | 0.087 | 0.122 | 0.109 |
| GO:0070628 | 0.23 | 0.424 | 0.081 | 0.111 | 0.1 |
| GO:0004198 | 0.07 | 0.076 | 0.054 | 0.076 | 0.069 |
| GO:0098632 | 0.25 | 0.465 | 0.087 | 0.122 | 0.109 |
| GO:0008375 | 0.087 | 0.094 | 0.068 | 0.094 | 0.085 |

**Supplementary Table S3.** Rate parameters used for growth models. Ranges are shown in square brackets, units are in hr^-1^.

| Subline<br>Parameter | PC9-DS1 | PC9-DS3 | PC9-DS4 | PC9-DS6 | PC9-DS7 | PC9-DS9 | PC9-DS8<br>(two-state) |
| --- | --- | --- | --- | --- | --- | --- | --- |
| $k_{\text{div,pre-drug}}$ | $0.04 * \ln(2)$ | $0.04 * \ln(2)$ | $0.04 * \ln(2)$ | $0.04 * \ln(2)$ | $0.04 * \ln(2)$ | $0.04 * \ln(2)$ | $0.04 * \ln(2)$ |
| $k_{\text{dth,pre-drug}}$ | $0.005 * \ln(2)$ | $0.005 * \ln(2)$ | $0.005 * \ln(2)$ | $0.005 * \ln(2)$ | $0.005 * \ln(2)$ | $0.005 * \ln(2)$ | $0.005 * \ln(2)$ |
| $k_{\text{div1,post-drug}}$ | $[0.022, 0.060] * \ln(2)$ | $[0.018, 0.060] * \ln(2)$ | $[0.023, 0.060] * \ln(2)$ | $[0.018, 0.060] * \ln(2)$ | $[0.018, 0.060] * \ln(2)$ | $[0.018, 0.060] * \ln(2)$ | $[0.021, 0.04] * \ln(2)$ |
| $k_{\text{dth1,post-drug}}$ | $[0.0227, 0.0625] * \ln(2)$ | $[0.0155, 0.0598] * \ln(2)$ | $[0.0259, 0.0653] * \ln(2)$ | $[0.0173, 0.0613] * \ln(2)$ | $[0.0196, 0.0630] * \ln(2)$ | $[0.0185, 0.0621] * \ln(2)$ | $[0.0205, 0.0615] * \ln(2)$ |
| $k_{\text{div2,post-drug}}$ | N/A | N/A | N/A | N/A | N/A | N/A | $[0.021, 0.040] * \ln(2)$ |
| $k_{\text{dth2,post-drug}}$ | N/A | N/A | N/A | N/A | N/A | N/A | $[0.0255, 0.0675] * \ln(2)$ |

### SUPPLEMENTARY FIGURE LEGENDS

**Supplementary Figure S1.**
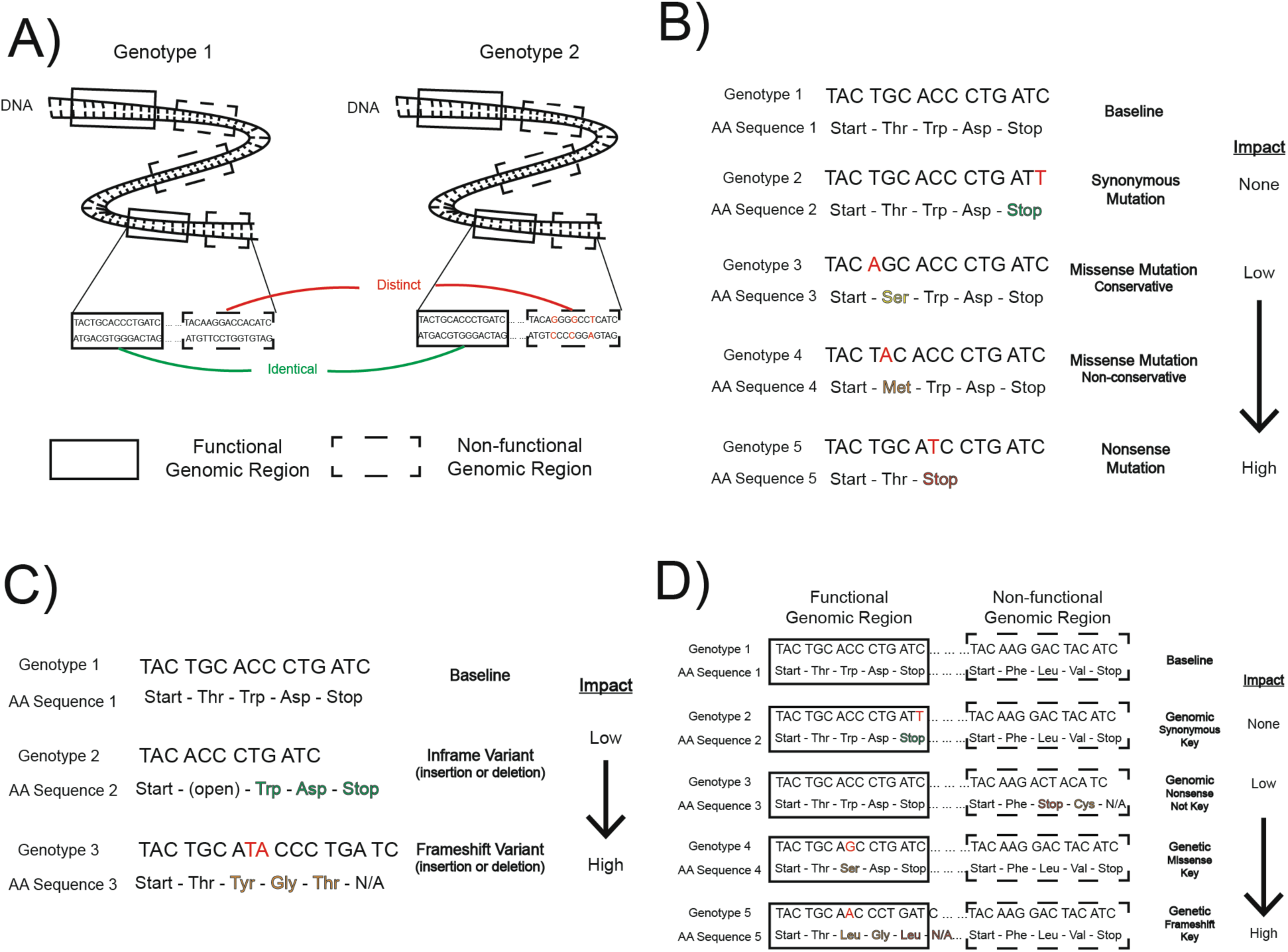
Clarification on the impact of mutational events on phenotype. (A) Cartoon illustration of two genomes that are genomically distinct but genetically identical, with respect to phenotype (i.e. drug-response sensitivity). In this illustration, functional genomic regions are highlighted with a solid black box, while non-functional regions are denoted with a dotted line. (B) Highlighted genomic region (from *A*) with substitutions, specifically single nucleotide variants (SNVs, most common variant type), and their resulting consequences. Nucleotide mutations are highlighted in red, and their resulting amino acid changes are highlighted based on the expected level of impact (green = no impact, yellow = minimal impact; orange = moderate impact; red = high impact). An impact column is provided as well, representing the same information. (C) Highlighted genomic region (from *A,B*) with insertion/deletion (indel) variants and their resulting consequences. Nucleotide insertions are noted in red, while deletions are shown with a shorter ‘genotype’ and open region of the amino acid sequence. Amino acid changes are colored as in *B*. Example indel types (inframe or frameshift) are shown as examples, but each type can lead to different levels of impact depending on the change to the underlying amino acid sequence. An impact column is provided for clarification (D) Example categorization of mutations with varying context (i.e., functional genomic regions), classification (i.e., substitution or indel), and consequence (i.e., synonymous, nonsynonymous – missense, nonsense). Functional genomic regions and non-functional regions (same as *A*) are highlighted in the same genotype. Nucleotide mutations, amino acid changes, and impact are noted in the same way as *B* and *C*. The combination of context, classification, and consequence define whether a mutation falls into the genomic or genetic category, noted in the figure.

**Supplementary Figure S2.**
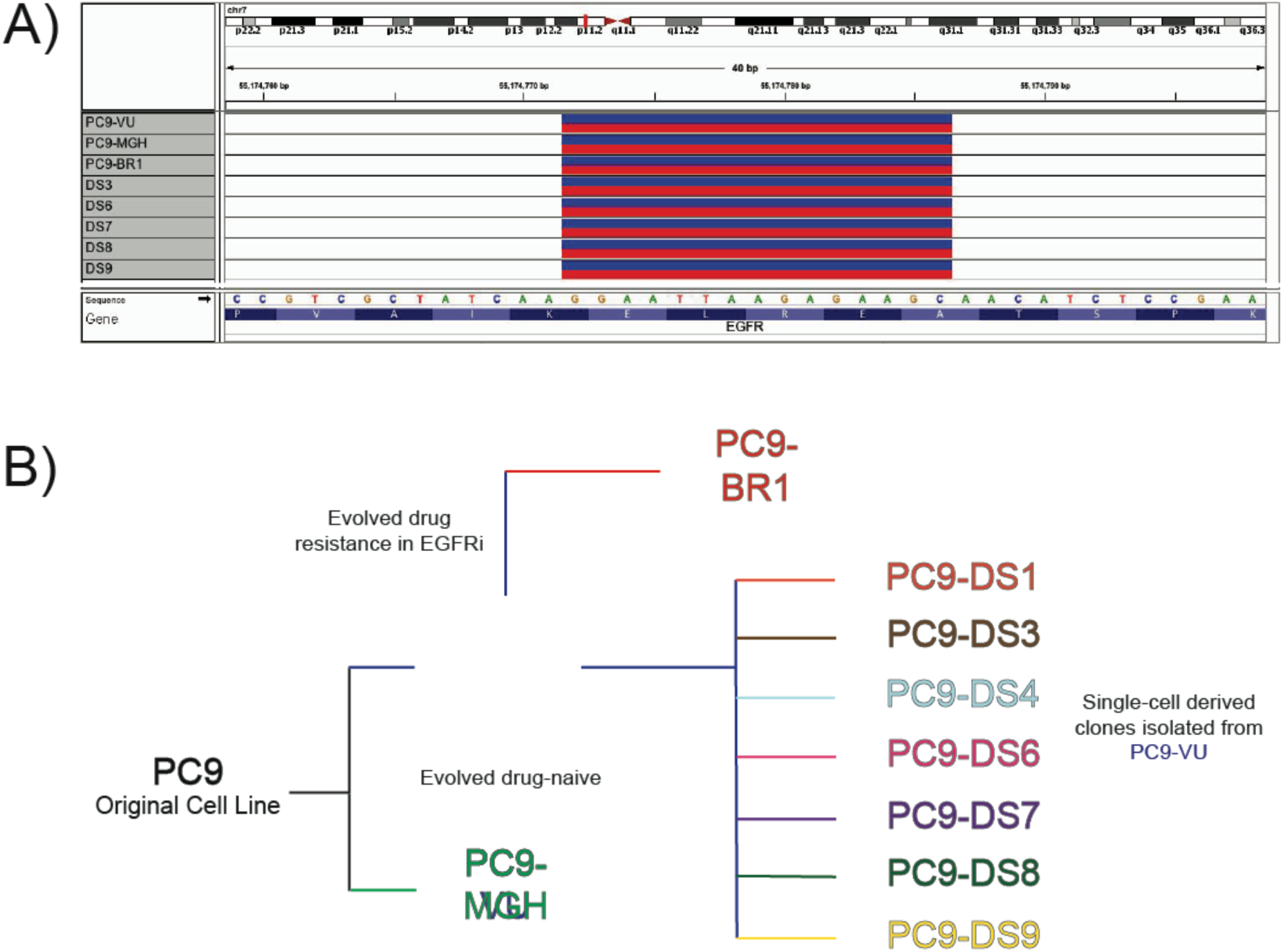
Understanding the PC9 Cell Line Family. (A) Identification of canonical EGFR-ex19del in PC9 cell line family members. As a positive control, we detected the same exon 19 deletion (common in EGFR-mutant non-small cell lung cancer (NSCLC)) mutation. Plotted as a screenshot from the Integrative Genomics Viewer (IGV). Red corresponds to potential deletions and blue corresponds to potential insertions. (B) Schematic representation of PC9 cell line family. Two ‘versions’ of NSCLC cell line PC9 were separately maintained in culture without selective pressure at two different universities (VU – Vanderbilt University; MGH – Massachusetts General Hospital). A resistant cell line was derived in EGFR inhibition (EGFRi, afatinib) from PC9-VU, named PC9-BR1. Several discrete sublines (DS*n*) were single-cell isolated from the PC9-VU cell line version. Together, these cell line versions and sublines make up the PC9 cell line family. Coloration is consistent with data visualizations in main and supplementary information.

**Supplementary Figure S3.**
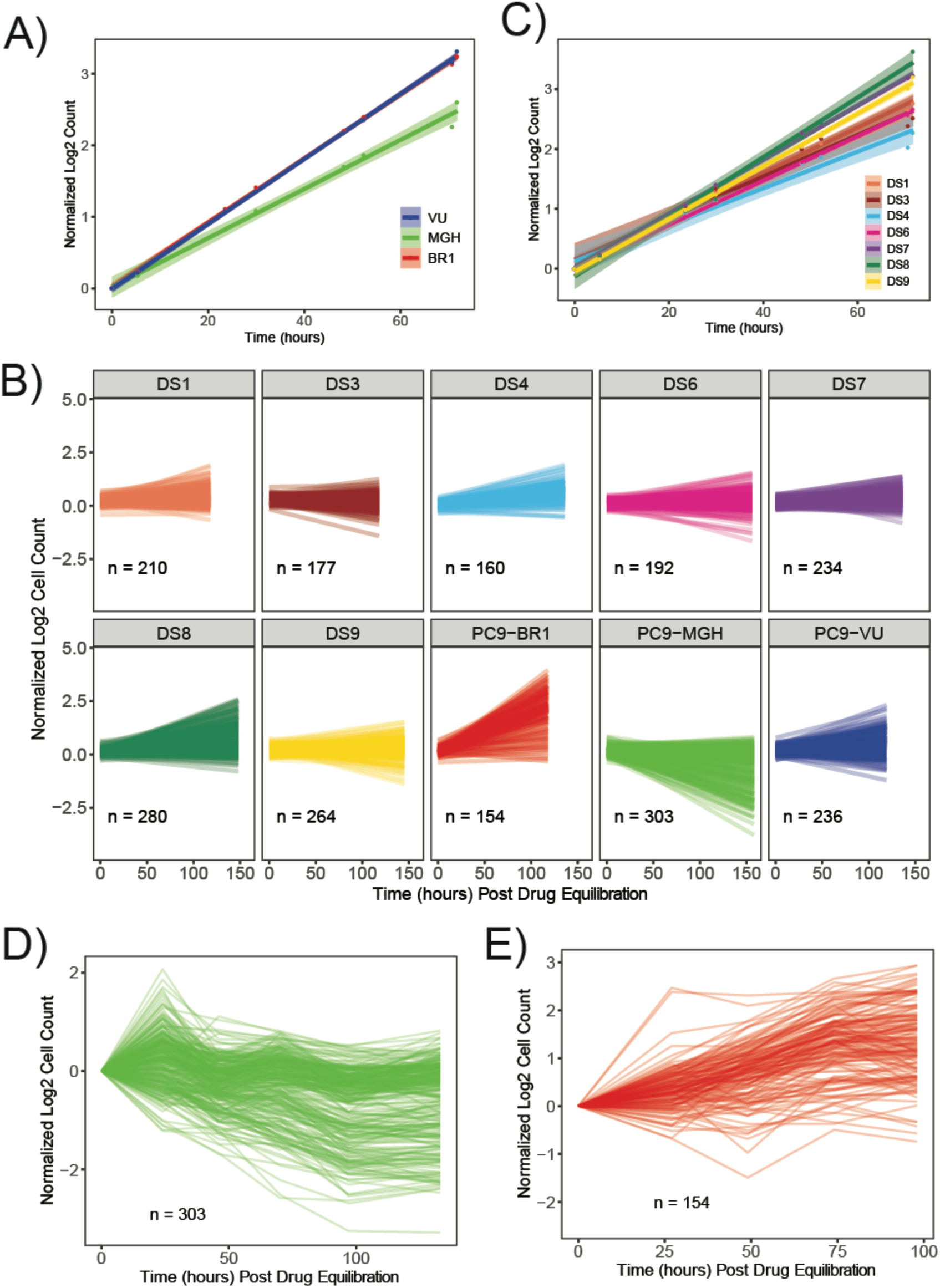
Visualizations of drug-naïve DIP rate assays and cFP trajectories. (A) Cell proliferation trajectory of cell line versions with vehicle (DMSO) treatment. Versions all grow exponentially, with PC9-VU and PC9-BR1 growing at the exact same rate. Dots represent means of six experimental replicates with a linear model fit and 95% confidence interval. (B) Linear model fits of clonal fractional proliferation (cFP) assay trajectories for each experimentally tested cell line version and subline in EGFRi (Erlotinib). Number of colonies (n) in each assay are noted within plot. Model fits were normalized to ∼48h post drug treatment. (C) Cell proliferation trajectory of discrete sublines (single-cell derived clones of PC9-VU) in vehicle treatment. Sublines grow at nearly the same exponential growth rate. Dots represent means of six experimental replicates with a linear model fit and 95% confidence interval. (D) cFP trajectories for PC9-MGH cell line version in EGFRi. Data are normalized to approximately 72 hours post EGFRi treatment. (E) cFP trajectories for PC9-BR1 cell line version in EGFRi. Data are normalized to approximately 72 hours post EGFRi treatment. Experiment was stopped after multiple colonies reached confluency.

**Supplementary Figure S4.**
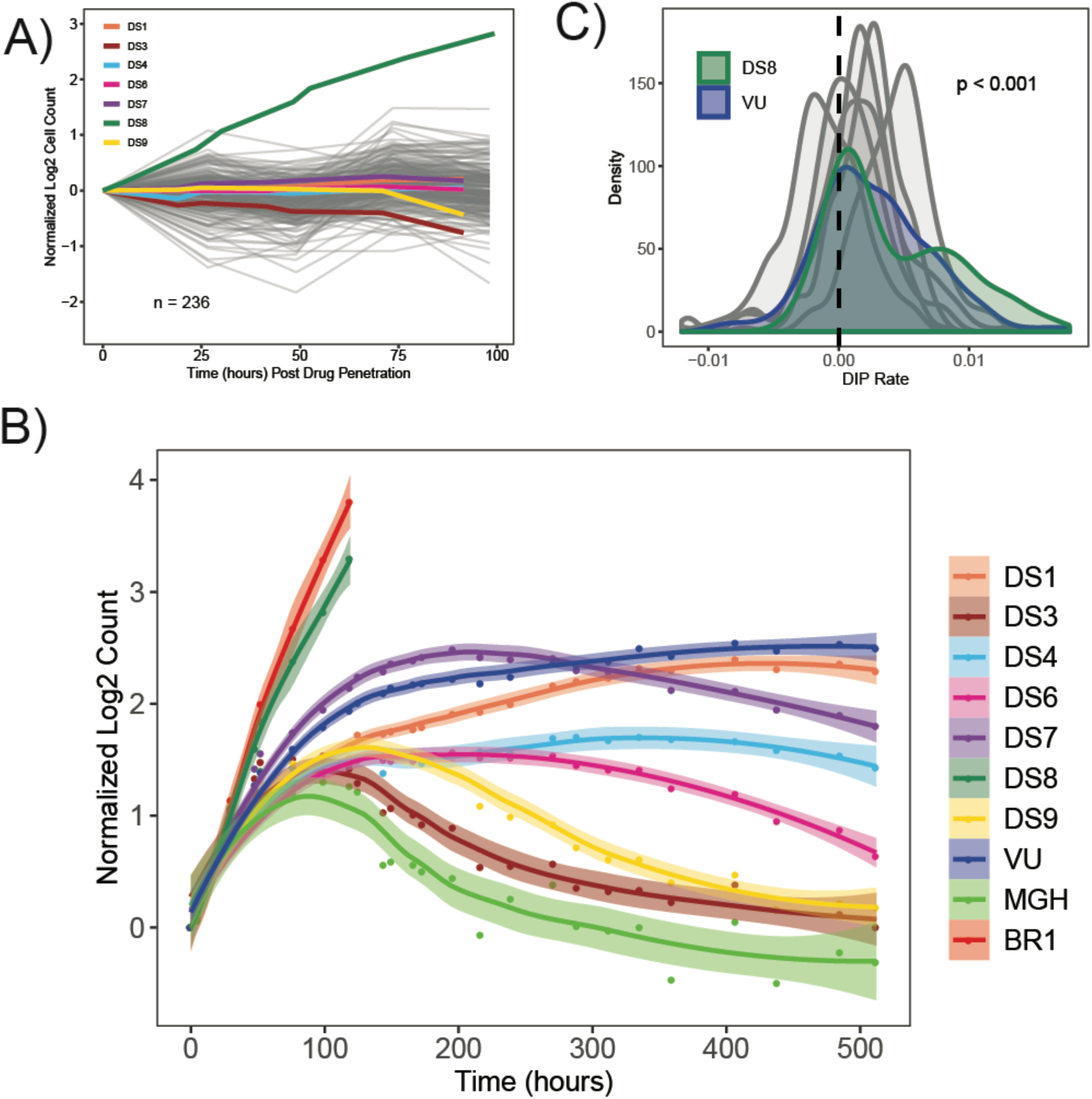
Direct comparison of cell line versions and sublines drug response profiles. (A) cFP trajectories for PC9-VU cell line version (grey) compared to population-level responses of PC9-VU sublines (colors) in EGFRi (Erlotinib). cFP trajectories are normalized to approximately 72 hours post EGFRi treatment. Sublines are normalized to approximately 125 hours post EGFRi treatment, with the exception of DS8. DS8 was normalized to the time of treatment (time = 0, reached confluency during the course of experiment because resistant to EGFRi). Means of subline time point replicates are plotted. (B) Full population-level drug-response trajectories of cell line versions and sublines in EGFRi. Data are normalized to the time of treatment. The loess linear fit is shown through time point replicate means (points), with a 95% confidence interval. (C) cFP DIP rate distributions for PC9-VU (blue) and sublines (DS1, DS3, DS4, DS6, DS7, DS9 = grey; DS8 = seagreen). A Mood’s median test was performed between all sublines, and medians were found to be statistically significant (p<0.001).

**Supplementary Figure S5.**
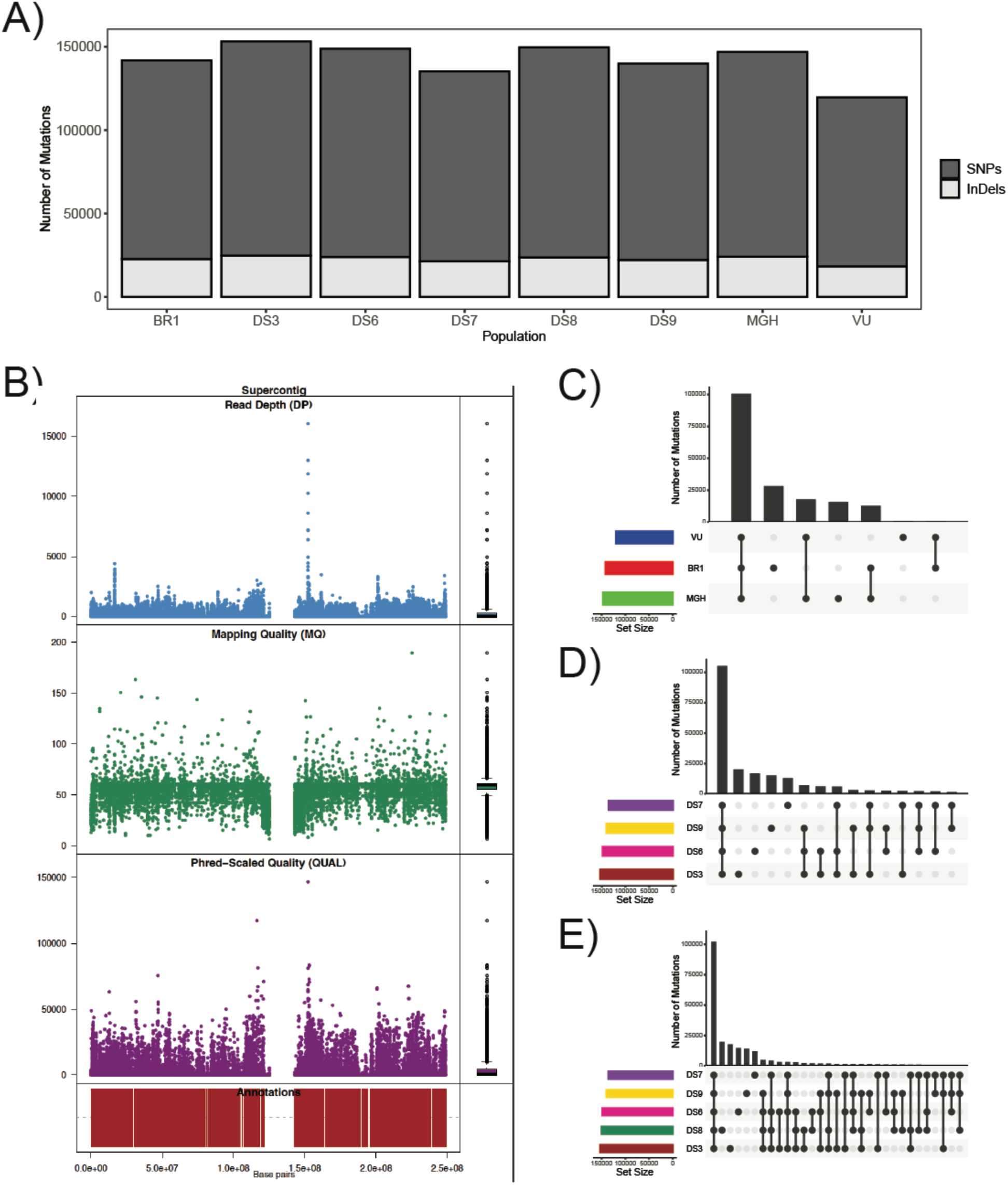
Metrics of whole exome sequencing data comparison across PC9 cell line family. (A) Total number of mutations identified through variant calling compared to hg38 reference genome. Mutations are separated into substitutions, specifically single-nucleotide polymorphisms (SNPs), and insertions and deletions (InDels). (B) Sequencing quality metrics for PC9 cell line family (considered together as one group). Read depth (DP) is a measure of sequence coverage; Mapping quality (MQ) details how well the sequencing reads are mapped to the reference genome; Quality (QUAL) is a score developed for Phred base calling that measures the confidence in called variants based on sequencing error probabilities; Variant count is a reflection of the variants per site identified over small sections (windows) of the reference genome. (C) Quantified Venn diagram (i.e., UpSet plot) of unique, and intersections of, mutations in cell line versions. (D) UpSet plot of unique, and intersections of, mutations in sublines, excluding DS8. (E) UpSet plot of unique, and intersections of, mutations in sublines, including DS8. Unique mutations in *C-E* are also represented in FIGS. 3B, 4B, and 6A.

**Supplementary Figure S6.**
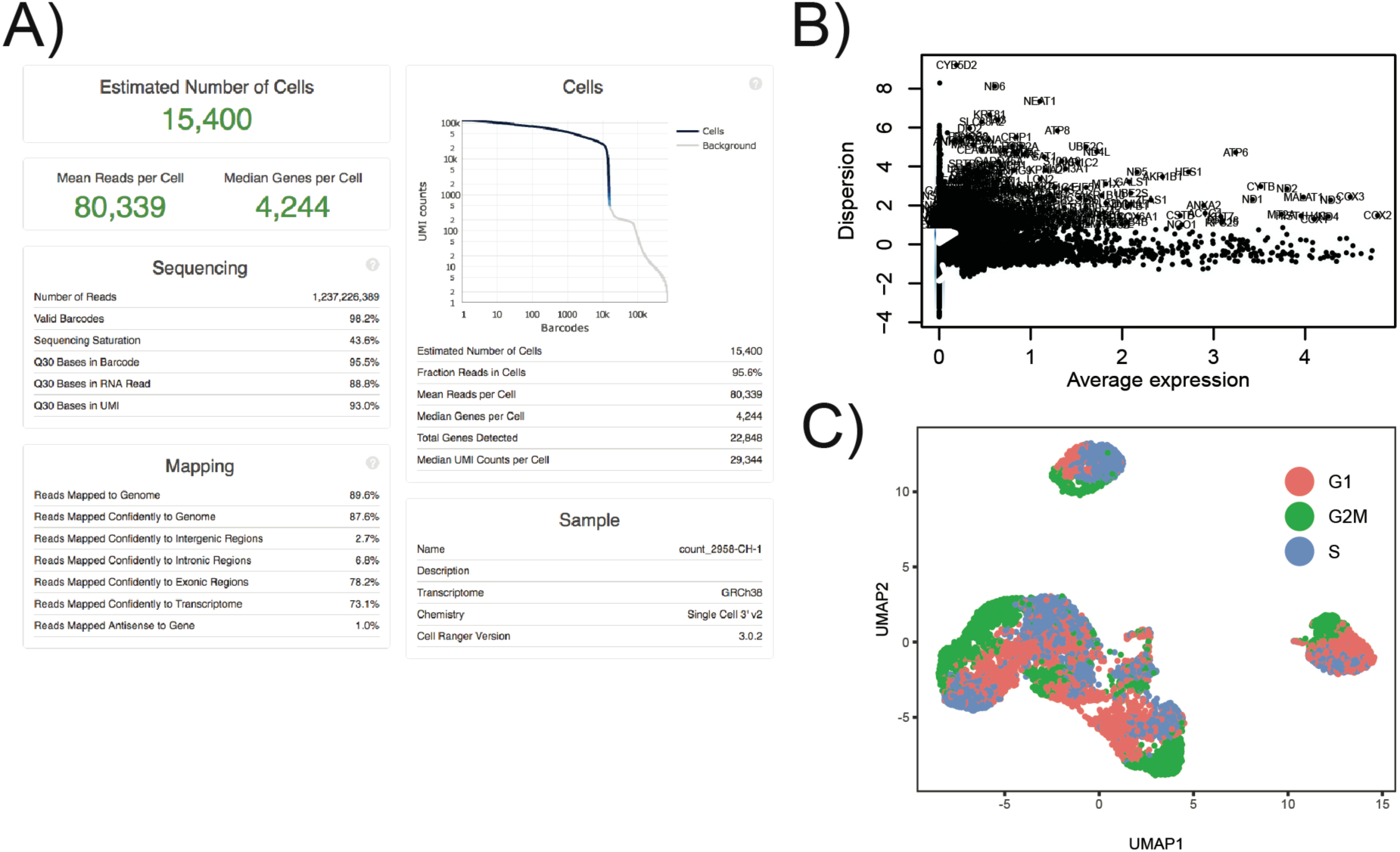
Metrics for sc-RNA-seq comparison across PC9 cell line family. (A) Cell Ranger (i.e., computational tool for generating count matrix) output file detailing metrics of sequencing run (quality, mapping, barcode identification, etc.). (B) Feature identification for genes that transcriptomically differentiate PC9 cell line family. Variable genes are projected on a plot of dispersion vs. average gene expression, and gene names that pass feature selection threshold are noted (0.1 < average gene expression < 8, log variance-to-mean ratio > 1; 499 genes). (C) UMAP projection of PC9 cell line family (see same projection in Supplementary FIG. S7), colored by cell cycle score. No discernible separation was identified between members of the family on this scale, and therefore regressed data was not used for further analyses.

**Supplementary Figure S7.**
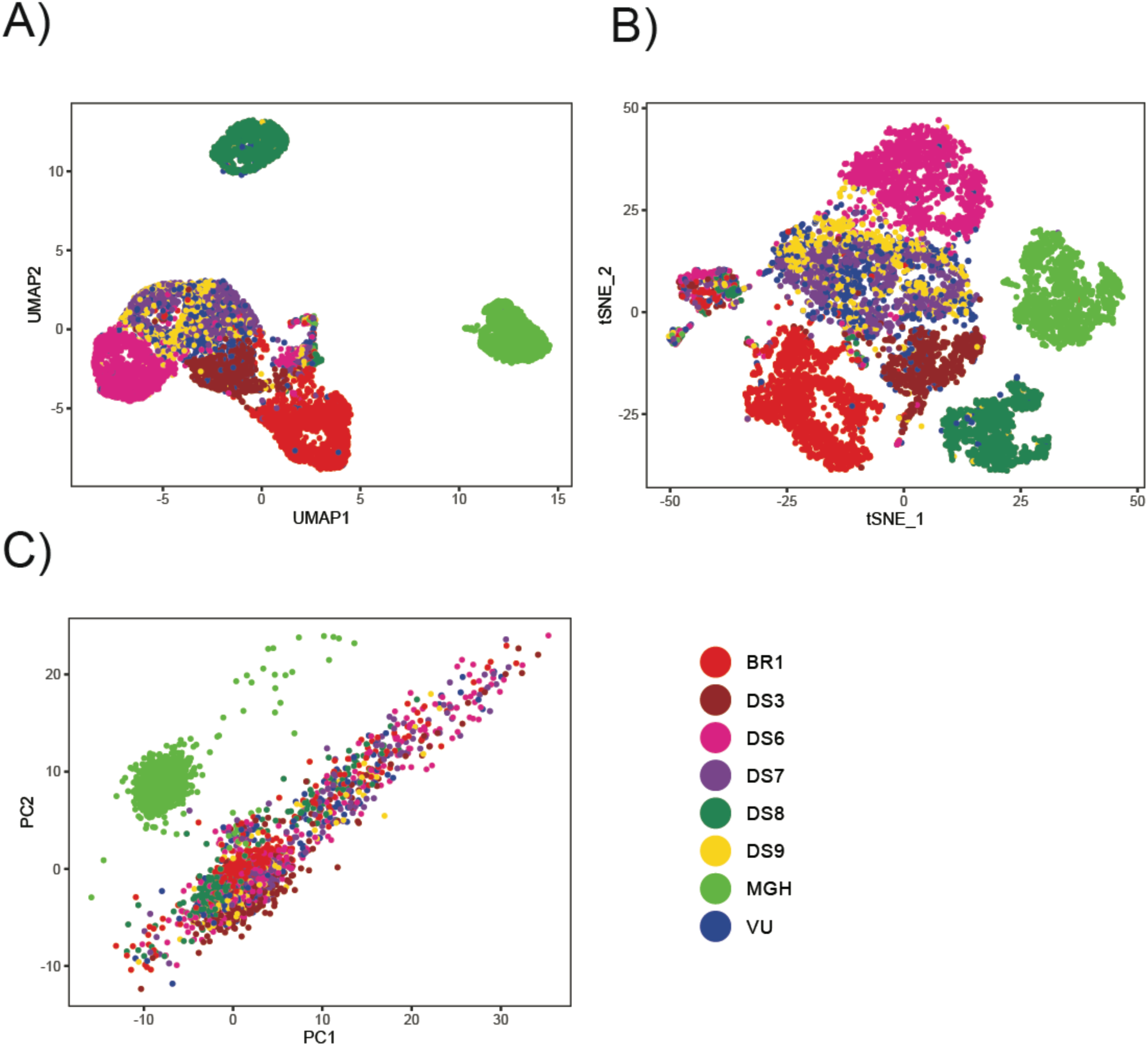
Additional sc-RNA-seq visualizations of PC9 cell line family using different dimensionality reduction approaches. (A) UMAP projection of PC9 cell line family, colored by family member. This visualization is the same space visualized in FIG. 3G, 4G, and 6C of the main text, but with all PC9 family members (cell line versions and sublines) included. (B) t-distributed Stochastic Neighbor Embedding (t-SNE) representation of sc-RNA-seq data. Similar separation and overlap can be seen as in the UMAP projection, but relative differences are not as discernible. (C) PCA representation of sc-RNA-seq data. This projection shows the clear transcriptomic differences between PC9-MGH and cell populations derived from PC9-VU.

**Supplementary Figure S8.**
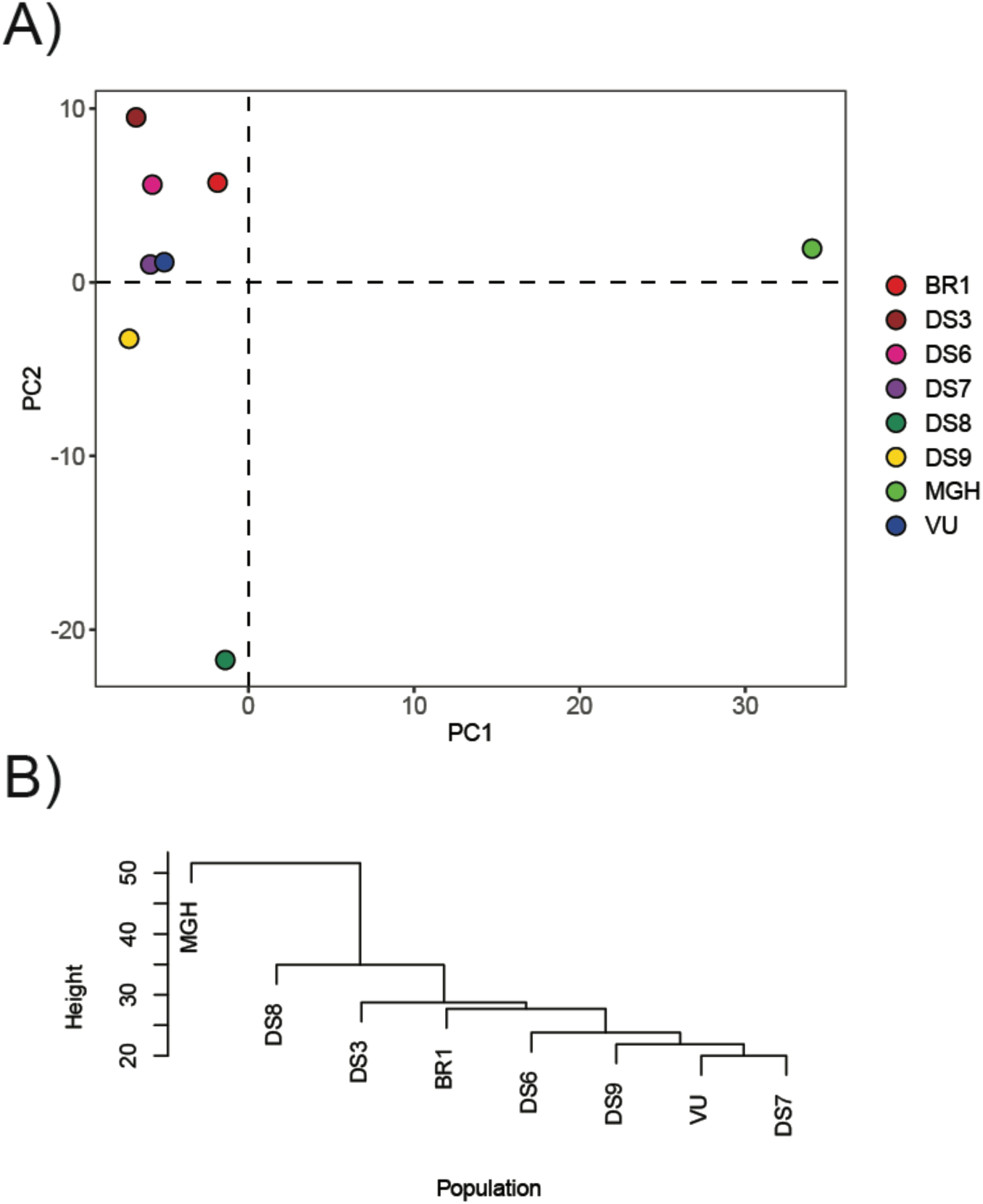
Confirmatory bulk RNA sequencing of PC9 cell line family. RNA sequencing (RNA-seq) was used to confirm sc-RNA-seq transcriptomic phenotypes. (A) PCA of single-replicate normalized RNA-seq count data. Similar distinctions and distances exist between this reduction and the UMAP projection in FIG. S7A. (B) Hierarchical clustering of RNAseq normalized count data. Clustering was performed on the pairwise Euclidian distance matrix created from the relative log transformed gene counts using the Ward’s minimum variance method. The clustering is consistent with the distances seen in *A*.

**Supplementary Figure S9.**
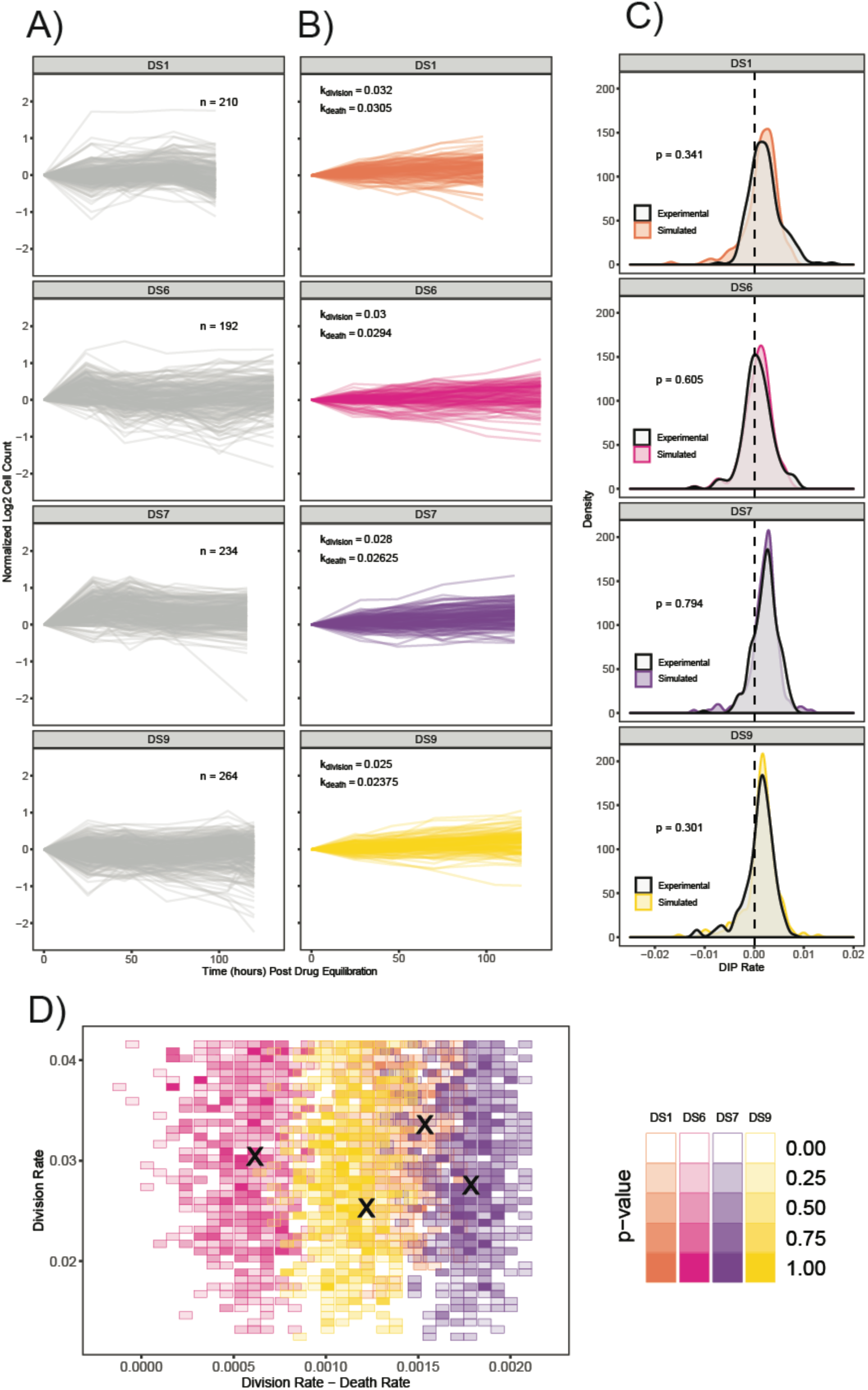
Monoclonal growth model (MGM) simulations and comparison to experimental cFP data. (A) Experimental cFP time courses for additional PC9-VU sublines (DS1, DS6, DS7, and DS9) in response to 3μM Erlotinib (trajectories corresponding to DIP rate distributions in FIG. 2D). Each trace corresponds to a single colony, normalized to 72 hours post-drug treatment. Only colonies with cell counts greater than 50 at the time of treatment were kept. (B) *In silico* cFP time courses with division and death rate constants that closely reproduce the experimental time courses in *A*. Trajectories are normalized to the time at which the simulated drug treatment was initiated, and simulated cell counts are plotted only at experimental time points. Although the simulation was initiated with the same number of trajectories as colonies in the corresponding experiment (for that subline, *n*), only colonies with initial cell counts greater than 50 at the time of simulated drug treatment are shown. (C) Comparison of experimental and simulated DIP rate distributions calculated from time courses in *A* and *B*. Distributions are compared statistically using the Kolmogorov-Smirnov (K-S) test (see Methods). A dashed black line signifies zero DIP rate, for visual orientation. (D) Parameter scan of division and death rate constants for the four sublines in *A*–*C*. For each pair of rate constants, we ran model simulations (same number as corresponding subline), calculated DIP rates and compiled them into a distribution, and then statistically compared against the corresponding experimental DIP rate distribution using the K-S test. Color shades correspond to p-values; all p<0.1 are colored white, indicating lack of statistical correspondence to experiment. **×** denotes a division and death rate constant used in *B*.

**Supplementary Figure S10.**
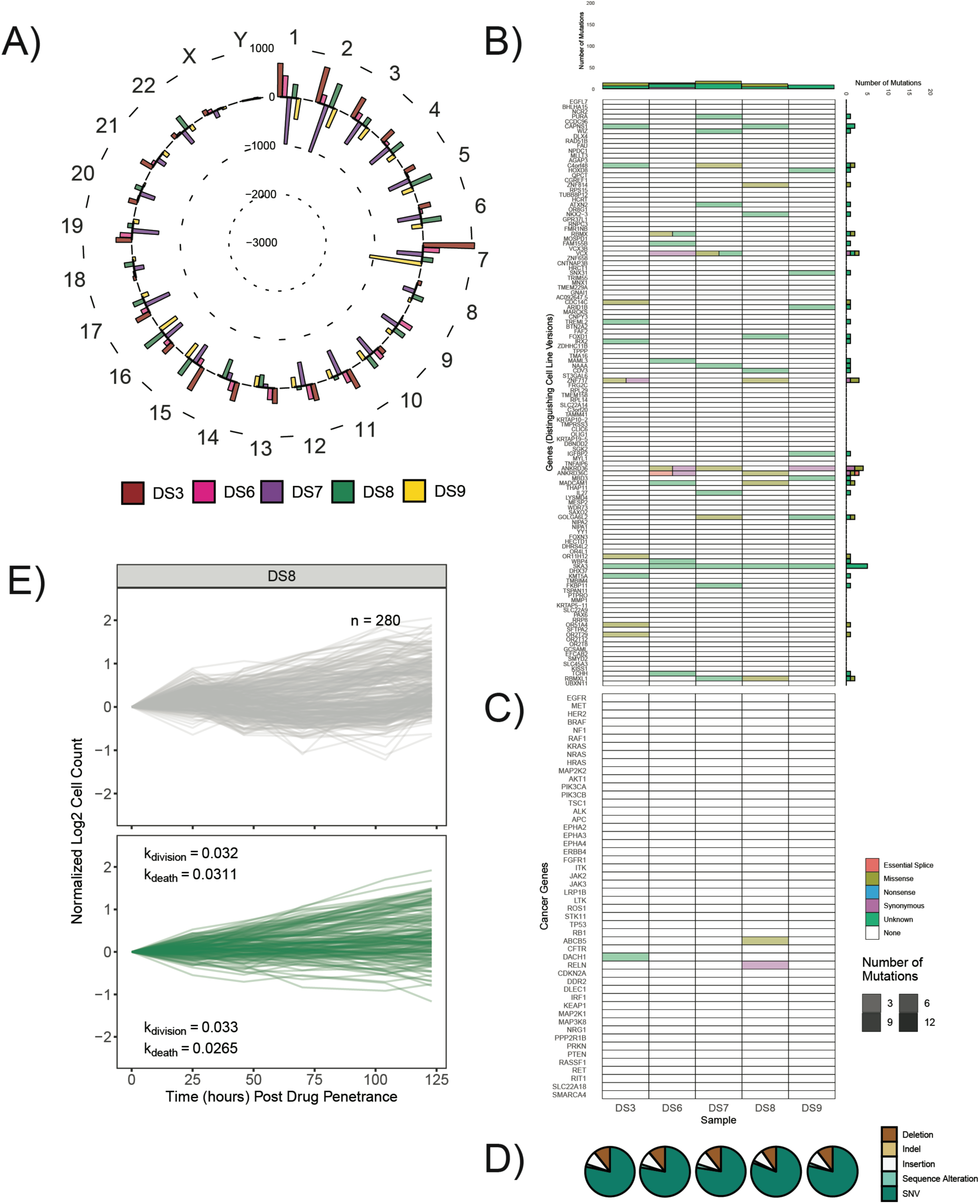
Additional visualizations of DS8 genomic, drug-response, and model-simulated data. (A) Mean-centered mutation count by chromosome for all sublines, including DS8. For each chromosome, sublines with fewer mutations than the mean have a bar pointing inwards, while those with more mutations point outwards. Chromosome numbers are noted on the outside edge of the circle. (B) Quantification of mutational differences between sublines, including DS8, based on a signature of genes with a high nonsynonymous mutational load in cell line versions. Heatmap elements are colored based on type of mutation: *Essential Splice Site*, *Missense*, *Nonsense*, *Synonymous*, and *Unknown*. Transparency of heatmap elements corresponds to the number of mutations in the gene-mutation type pair (more mutations, less transparent). Total numbers of mutations (stratified by mutation type) across genes and cell line versions are shown as bar plots to the right and above of the heatmap, respectively. (C) Same as *B*, but for a literature-curated set of cancer-associated genes implicated in lung cancer. (D) Mutation class pie charts. All unique mutations in each cell line version were classified as single nucleotide polymorphisms (SNPs), insertions, deletions, indels (combinations of insertions and deletions), or sequence alterations. (E) Example experimental (top) and simulated (bottom) cFP time courses for DS8, using the same scheme as *Supplementary* FIG. S9. Two sets of division-death rate pairs are included in the simulated data, to represent the two states.

**Supplementary Figure S11.**
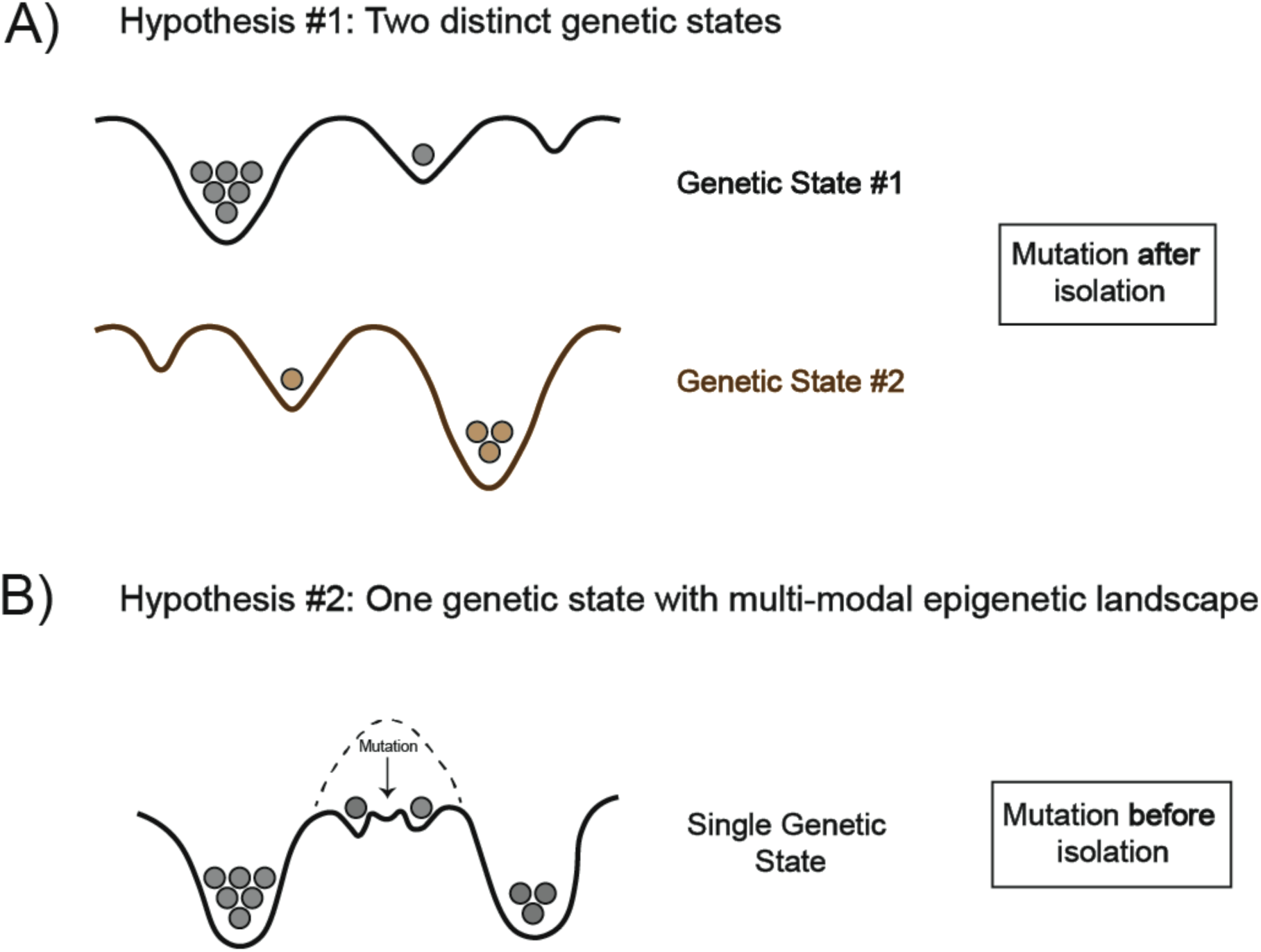
Potential explanations for DS8 genetic variability. DS8 exhibits extensive genetic, transcriptomic, and drug response differences compared to other members of the PC9 cell line family. DS8 also exhibits heterogeneity in the sc-RNA-seq (representation in different regions of UMAP projection) and cFP data (bimodal DIP rate distribution), which are the assays that can identify heterogeneity at the single-cell level. Therefore, this heterogeneity corresponds to at least one new genetic state. According to the G/E/S heterogeneity framework, this heterogeneity could come from the following two sources: (A) DS8 is composed of two distinct genetic states, each with their own epigenetic landscapes. In this explanation, the left mode of the DS8 DIP rate distribution corresponds to remnants of a subline isolated from the a single epigenetic basin associated with the PC9-VU genetic state. The right mode of the distribution corresponds to a new genetic state, also with its own epigenetic landscape. Each of these genetic states may have multiple epigenetic basins occupied at biased, but potentially varying proportions (see also FIG. 7). Given this hypothesis, the mutation would have happened after single-cell isolation, in order to retain the PC9-VU epigenetic basin. (B) Alternatively, DS8 could be composed of a single genetic state with multiple basins heavily occupied in its epigenetic landscape. In this case, DS8 developed a genetic mutation while growing within the PC9-VU cell line version, and was isolated during single-cell cloning. The mutation could have modified the associated epigenetic landscape so as to lower the energy barrier between stable basins. In this case, the bimodal DIP rate distribution corresponds to two epigenetic basins of the same genetic state, as opposed to two distinct genetic states. Either way, DS8 incorporates a new genetic state.

**Supplementary Figure S12.**
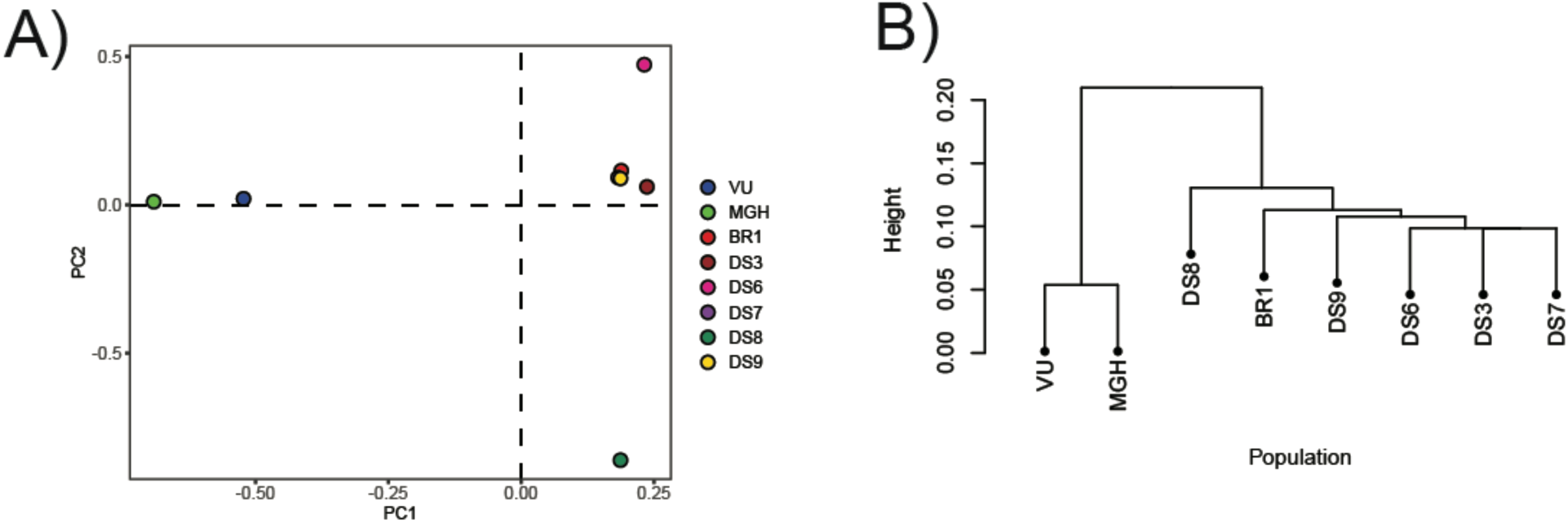
Genomic relatedness between PC9 cell line family members. (A) PCA of PC9 genotypes. Using a subset of single nucleotide polymorphisms (SNPs) in approximate linkage equilibrium, a genetic covariance matrix was calculated. The covariance matrix was converted to a correlation matrix to achieve appropriate scaling, and PCA was run to identify SNP eigenvectors (loadings of the principal components). (B) Hierarchical clustering of PC9 genotypes. Using an identity-by-state analysis, a matrix of genome-wide pairwise identities was calculated. Hierarchical clustering was performed on these identities to determine sample relatedness.

**Supplementary Figure S13.**
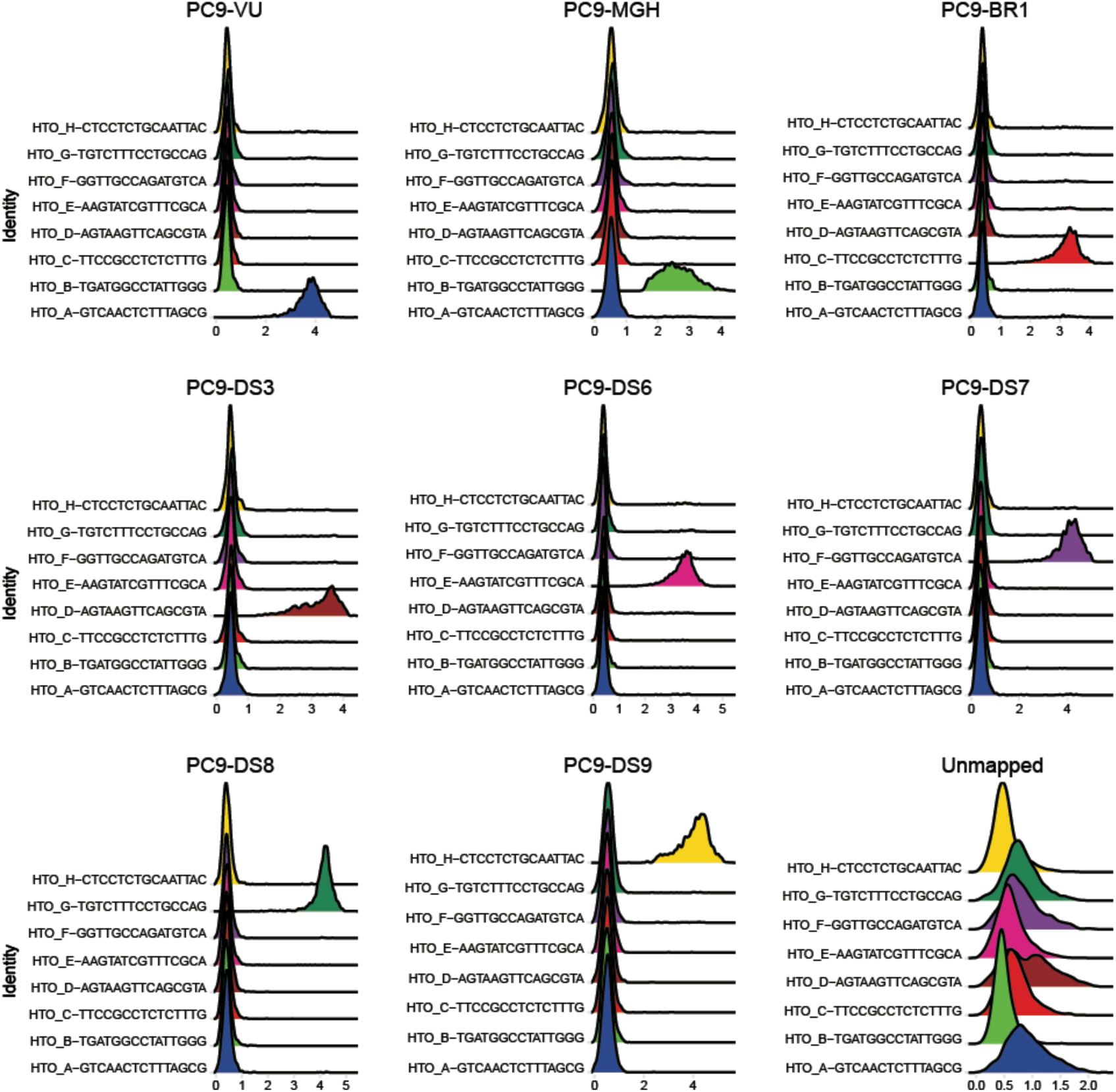
Sample identification in “hashed” PC9 cell line family members. Proportional representation of cell populations with each of eight specific “hashtag” antibodies based on the hashtag oligonucleotide (HTO) expression level. Each sample has a single corresponding HTO, while a minority of the HTOs reads were unmapped.

